# Time-resolved single-cell state tracing exposes bidirectional state dynamics during differentiation

**DOI:** 10.64898/2026.09.18.752672

**Authors:** E. Gonzalez Sanchez, F. Fabro, J. Klavert, G. Vroeg in de Wei, J. Boers, B. Tan, A. Sacchetti, M. van Leeuwen, E. Bindels, R. Boers, J. Gribnau, M. P. Creyghton

## Abstract

Differentiation dynamics have been challenging to study across time as measuring cell state, transcriptionally or epigenetically, typically requires its destruction. Here, we developed single-cell state tracing using the DCM-Time machine to enable the characterization of two transcriptomes, separated in time in single cells during epithelial cell differentiation in the intestine of living animals, and during neuronal differentiation in culture. We show that retrospective transcriptional histories and current states are consistent with bidirectional movements along differentiation trajectories both in vivo and in vitro that become rarer as cells mature. Furthermore, we observe rare transcriptome conversions, across canonical lineage boundaries. Our data support a model in which differentiating cells occasionally fail to stabilize lineage-associated programs, enabling reversals and subsequent resolution toward alternative lineage states.

## Introduction

During development, stem cells differentiate to give rise to a wide array of different cell types by activating cell type-specific gene expression programs that involve thousands of genes^1^. Activation of these programs typically occurs in a step wise manner with each step further restricting differentiation potential^2,3^, aligning with the classical view of differentiation as marbles rolling downhill. However, deviations of this linear model of differentiation, have been described^4^. These include cells reverting to a more stem cell-like state during damage and regeneration^5–7^, and the adoption of mixed lineage identities, a process that can be hijacked by cancer cells^4,8^. Nevertheless, some of these results remain difficult to interpret because lineage tracing typically relies on the activity of single marker genes^9^, which may be activated without formal differentiation (i.e. lineage priming)^10,11^ and thus be only partially specific as a proxy for full cell state.

The ability to temporally link transcriptional states across development is severely limited as the act of measuring the transcriptional or epigenomic state of a cell requires its destruction. This limits cell state measurements to single time points. To compensate for this lack of experimental insight, the analysis of cell state changes has primarily relied on in silico inference by logically linking related states following a time course of measurements ^12–21^. While these methods can provide important insight into the manner in which cells make lineage decisions, they lack true temporal coupling of states and rely on various sets of assumptions ^15^. Recently transcriptional history was measured in split cell cultures ^22^. However, this approach does not directly couple two timepoints in the same cells and is also incompatible with single cell analysis. Tracing of cell states across time has also been attempted by aspirating cytoplasm from cultured cells at multiple time points ^23^. While promising, this technique is laborious, low throughput and limited to cells in culture. Others have used Dam fusion proteins to trace chromatin state back in time ^24^. However, this method is limited to time intervals spanning hours rather than days. To overcome these limitations, we developed a single cell state tracing approach based on the DCM-Time machine^25^ to follow single cells back in time in living animals and cell cultures. Using DCM methylation as a retrospective proxy for prior transcriptional state, we find that cells can transiently activate lineage-associated programs before fully consolidating cell identity. These observations are consistent with multidirectional differentiation trajectories in both the mouse intestine and a neural differentiation model. Our analyses support a model in which cells that fail to stabilize an initial lineage program can remain plastic and, in rare cases, resolve toward alternative lineage states.

### Benchmarking single-cell MeD-Seq in mESCs using combinatorial indexing

To enable the measurement of two transcriptomes separated in time in single cells, we used the DCM-time machine mouse that we developed previously to trace cell state changes in bulk tissues^25^. The method relies on a bacterial methyl transferase (*dcm*) fused to the RNA polymerase II subunit B (*Polr2B*), which is primarily found in gene bodies of actively transcribed genes. Doxycycline (dox) inducible expression of this (*dcm-Polr2b*) fusion deposits a bacterial methylation mark on expressed genes (Fig. 1A), that is maintained in cells and further propagated at a relatively high efficiency in cycling cells^25^. Therefore, a controlled pulse of *dcm-Polr2b* expression provides a record of cell state that can be recovered at a later stage of development. To enable single cell state tracing, we set up a ligation-based combinatorial indexing approach that assigns a specific barcode combination to each nucleus drawing from several technologies^26–28^. This technology was rooted in MeD-Seq to enable efficient analysis on DNA methylation using the methylation sensitive enzyme LpnPI (Fig. S1A, sciMeD-Seq). Using this approach in mouse embryonic stem cells (mESCs) by low coverage sequencing (40,987 mean reads per cell), we recovered a number of unique barcode combinations that was similar to the number of input nuclei (Fig. S1B), demonstrating the robustness of this approach. Barcode collisions were calculated to occur at a rate of 2.26% and experimentally measured at 7.5% to include potential doublet formation (see methods). Metagene plots of sciMeD-Seq data in mESCs for combined single nuclei (pseudo-bulk) confirmed induction of DCM signal at gene bodies only after dox induction (Fig. S1C-F). Furthermore, we observed DCM induction at non-coding regions with active enhancer chromatin in mESCs and DCM depletion at heterochromatin marks compared to surrounding genomic regions (Fig. S1G, H) consistent with expected Pol2 occupancy. DCM methylation was present at pluripotency genes, (e.g. *Nanog*) and enhancers enriched for pluripotent transcription factor (TF) binding sites (Fig. S1I, J), similar to our bulk analysis^25^. This demonstrates that our single cell approach captures expected pluripotency-associated transcriptional-state features.

**Fig. 1.**
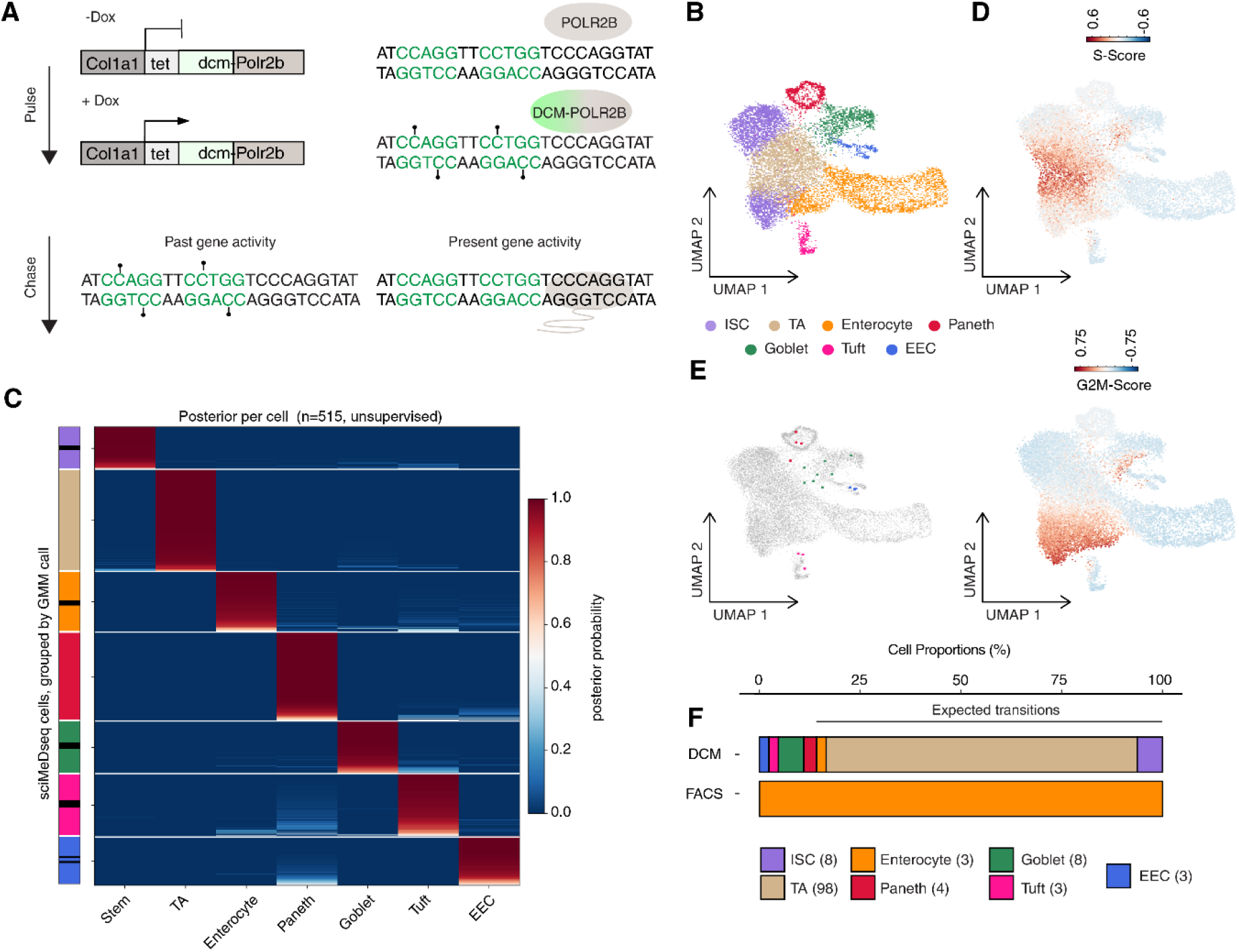
Combinatorial indexing based single cell state tracing of intestinal enterocytes. (A) Scheme of the Dcm-Polr2b cell tracing system. Upon dox induction, the DCM-POLR2B protein is generated, which methylates regions of ongoing transcriptional activity (pin marks). Following a chase period, past gene activity (DCM methylation) and current gene activity (mRNA) can be measured by MeD-Seq and RNA-Seq. (B) Integrated UMAP between scRNA-Seq reference data (Atlas and Large dataset (n = 7,216 and 10,396 cells, respectively))^31^ and sciMeD-Seq (n=515) datasets from FACS sorted enterocytes. The left UMAP represents the clusters corresponding to the main cell types of the small intestine while the right UMAP shows the reference cells colored in gray and DCM cells colored by their corresponding cell types (n=515 nuclei). (C) Posterior probability of cell- type assignment per single cell, based on a Gaussian Mixture Model (GMM) fit to DCM module Z-scores of cell- type-specific marker gene sets. Each row is DCM data for a single cell; each column is one of the 7 gene-sets corresponding to annotated cell types as indicated. Rows are ordered by assigned cell type. Color encodes posterior probability, from dark blue (0) to dark red (1). (D) UMAP showing cell cycle markers gene sets (S score, top; G2M score, bottom), and (E) highlighting sorted enterocytes with secretory-history profiles, colored by their corresponding cell type (right). (F) Transitions of cells with concordant labelling between GMM and BBKNN integration-based annotations. The top column shows cell labels based on DCM GMM and integration, and the bottom column shows FACS based origin (enterocytes).

### In vivo single-cell state tracing in the mouse intestine reveals secretory to absorptive state discordance

Next, in order to trace cell states back in time during in vivo differentiation, we used the mouse intestine as a model. During differentiation of the intestinal epithelium, intestinal stem cells (ISCs) either differentiate into secretory cells, including goblet, tuft, Paneth and enteroendocrine (EEC) cells, or they differentiate into enterocytes which function as absorptive cells^29,30^. The whole process takes approximately five days during which maturing cells migrate from the base of the crypt where the stem cells reside, towards the top of the villus where they are shed^29^. This is compatible with DCM propagation rates in the mouse intestine (calculated at 59% per cell division)^25^.

We fed mice water supplemented with dox for 48 hours and subsequently isolated cells from the small intestine (jejunum) followed by Fluorescence-Activated Cell Sorting (FACS) for mature enterocytes using Glut2 and EpCAM antibodies^25^. Single cell DCM methylation was measured using combinatorial indexing to trace the cell state of individual enterocytes back in time. We recovered 515 nuclei after quality filtering (Fig. S2A, mean 116,825 reads per cell). DCM methylation induction and the relationship between aligned reads and unique DCM recovery was more uniform across cells than to those for mESCs (Fig. S2B-D), R^2^ = 0.71 vs 0.97 in mESC and intestine, respectively).

To identify cell types based on DCM methylation, we first used batch balanced k nearest neighbors (BBKNN) integration of DCM methylation data with scRNAseq data (Fig. 1B, Fig. S2E, F) ^31^. As expected, we found that, following 48-hour dox induction, most enterocytes could be traced to either an enterocyte or an immature state (Fig. S2F). However, a large number of cells (n=63) had DCM methylation profiles resembling secretory-lineage transcriptional states. This is at odds with the typical differentiation trajectory for enterocytes, which under physiological conditions emerge from stem cells via a transient amplifying state (TA)^29^. To confirm this observation, we leveraged scRNAseq data^31^ to generate a set of cell type specific differentially expressed (DE) genes (logreg score >0.5, FDR < 0.05 (t-test)) including several known cell type specific marker genes (n=127 genes; Table S1) (Fig. 1B, S3A, B). We validated BBKNN based cell type annotations by showing that each annotated cell type exhibits significant enrichment of its corresponding gene-set module score (Fig. S3A). We then used these marker gene sets to assign DCM based cell histories by unsupervised Gaussian Mix Modeling (GMM) (Bayesian FDR < 0.05; Fig. 1C). Analysing only cells with labels strictly matching between integration and enrichment of these marker genes (n = 127 cells; Fig. S3B,C), we still observed secretory-like histories among current enterocyte-lineage cells (4 Paneth, 8 Goblet, 3 tuft and 3 EEC cells; Fig. 1D,E,F), representing 14.17% of all filtered transitions and 3.49% of input enterocytes. This frequency is higher than expected from FACS misclassification or barcode collisions (∼1% and 2%)^32^. Furthermore, these cells are unlikely to represent barcode collisions or doublets as this, would have resulted in mixed enterocyte identities. However, the exact frequencies of these transitions are difficult to establish due to high DCM drop-out rates. Furthermore, integration of the data with scRNA-Seq data suggests that some of these discordant histories may represent more immature secretory states reminiscent of multilineage priming^33,34^.

### Isolated processing of single cells reveals rare discordant lineage-associated state histories

To validate these results using an alternative method that controlled for confounders, (i.e. doublet formation, sorting ambiguities and ambient fragments), we designed a second single cell state tracing strategy (scMeD&RNA-Seq) drawing from plate-based approaches, such as DAM-ID&T and scNMT^12,35^. This approach involves reaction miniaturization and robotics to measure DNA methylation and mRNA expression in the same cell, with each cell being processed separately to avoid cross contaminations by ambient nucleotide fragments. To enrich for underrepresented cell types in the intestine, and make sure each well contained a single viable cell, we used image-based FACS (see methods and Fig. S4B). Single cell content was visually verified for each well to make sure no doublets were present. This enabled direct coupling of current transcriptomes with retrospective DCM-derived transcriptional-state histories in single cells. (Fig. S4A). Benchmarking the analysis, we detected a mean of 1,363 genes by mRNA expression across 997 cells, and 1,154 genes based on DCM methylation and an additional 24,348 CpG sites per cell across 657 high quality cells (Fig. S4C and D, mean 519,033 reads sequenced per cell). After pairing, 528 cells were considered high quality for both RNA and DCM signal and were retained for subsequent analysis.

As expected, feeding the mice dox for 48 hours increased DCM methylation in isolated intestinal cells over non-induced mice, which gradually declined following a 72-hour chase (Fig. 2A, Fig. S4E). BBKNN integration of scMeD&RNA-Seq data (both plate RNA (pRNA) and plate DCM (pDCM)) with the reference scRNA-Seq datasets was followed by leiden clustering to assign RNA based cell states (Fig. 2B, Fig. S4F, G). To assign high-confidence cell-type labels based on DCM methylation, independently of integration, we used a larger cell type–enriched gene set based on scRNA-seq cell clusters (log2FC >1; FDR < 0.0001; t-test n=3,973) followed by computing the normalized average expression scores for each cell per gene set (Methods, Table S1). We then applied an unsupervised Gaussian mixture modeling (GMM) approach. For each gene set, scores across all cells were modeled using a two-component GMM (i.e. low relative expression versus high relative expression), and cells were probabilistically classified based on posterior membership probabilities (Posterior probability >0.85, Bayesian FDR<0.05, Fig. 2C). Using this approach, we identified all major intestinal cell types including secretory and absorptive lineages across assays (Fig. 2D, S5A) and expected cell type specific marker genes were correctly enriched in labeled cells (Fig. 2E). Several cells (n=119, 18.11% of all filtered DCM cells) received two labels (mixed states) or more and these were typically a mix of neighboring cell states confirming that we are capturing known lineage trajectories (Fig. S5E, Fig. 2F-H, mixed states indicated by *). These cells were predominantly found to be integrated in the ISC / TA compartment and as such may represent more immature states in the process of lineage specification or cells that are lineage primed ^34^. By analyzing DCM to RNA transitions from cells that were confidently assigned to a canonical cell state by GMM (n=312, FDR<0.05), we observed an increase in stem cell / TA-like DCM histories for all major absorptive and secretory lineages in the pulse chase compared to the pulse alone (Fig. S5B, C) which is consistent with their temporary presence in the villus. Furthermore, as expected by the reduced time to transition, mature states were more stable in the pulse dataset compared to the chase (Fig. S5D). Expected lineage relationships captured as ISC- or TA-like DCM histories in cells were overall assigned to current absorptive or secretory states. (Fig. 2F and Fig. S5F).

**Fig. 2.**
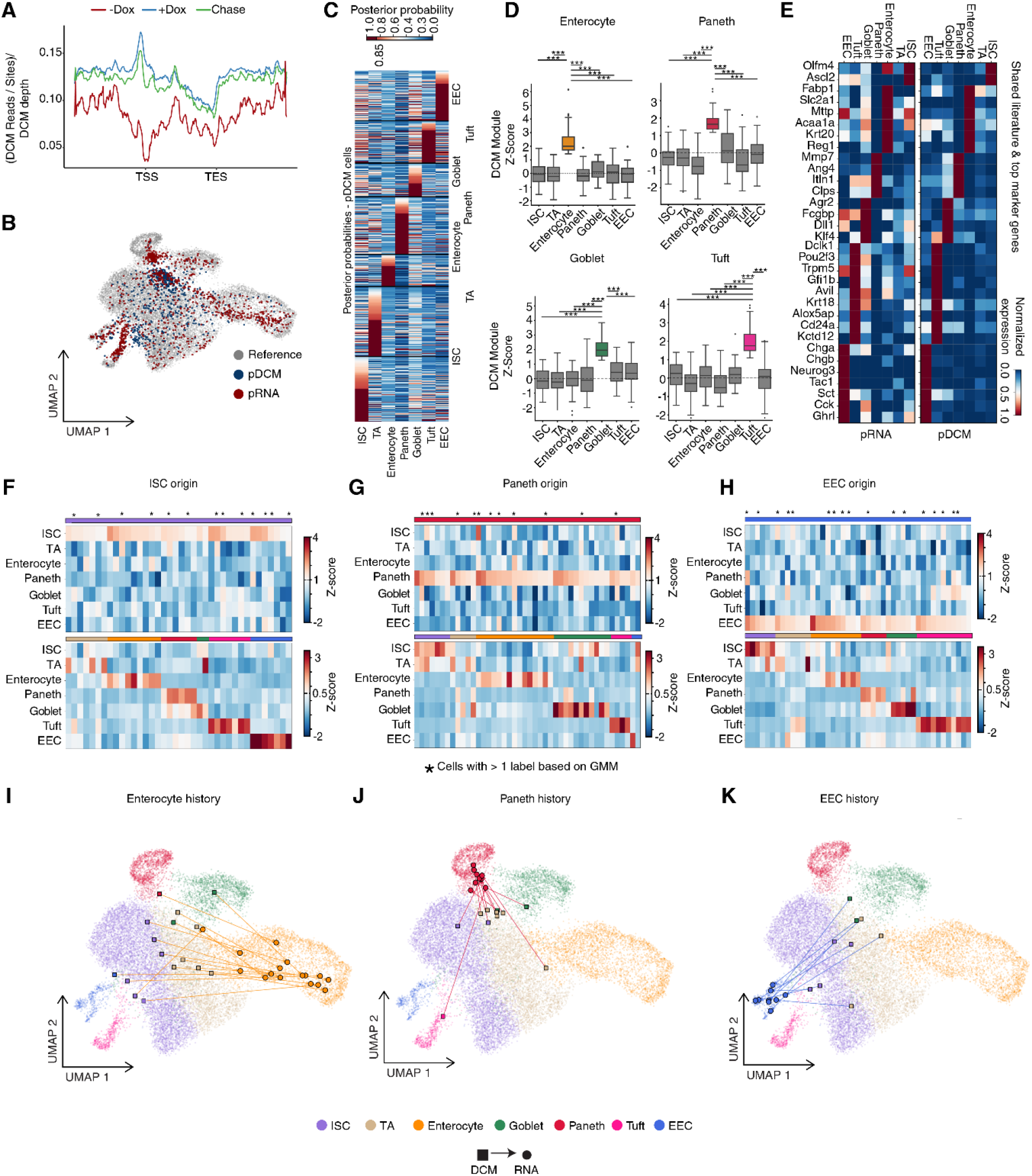
scMeD&RNA-Seq based single cell state tracing in the mouse intestine. (A) Metagene plot with the distribution of DCM reads for each condition (non-induced (-Dox), 48h dox (+Dox) and 48h dox followed by 72h chase (Chase)) across genes normalized for DCM sites and sequencing depth. (B) Integrated UMAP of reference scRNA-Seq data (grey) ^31^ and plate based scRNA-Seq data (red) as well as plate-based scDCM data (blue; left panel). (C) Heatmap showing module score based posterior probabilities generated using Gaussian Mix Modelling (GMM, >0.85 posterior probability). (D) DCM module Z-scores for marker gene sets specific to each cluster (cell type; ISC, n=603 genes; TA, n=295; Enterocyte, n=1,275; Paneth, n=123; Goblet, n=465; Tuft, n=696; EEC, n=475) calculated for indicated cell types following GMM-based cell assignments (FDR<0.05). Each panel indicates a separate cell type specific gene module tested. Z-score indicates deviation from the average expression pattern across all cells in the dataset. Values are compared using a pairwise t-test adjusted for multiple testing (FDR, *** (p < 0.001)). (E) Heatmap plot visualizing marker gene expression across cells for each cluster, shown separately for plate-based RNA data (left panel) and plate-based DCM data (pDCM; right panel). Color indicates depth-normalized and log-transformed gene expression levels. (F-H) pDCM and pRNA data scaled across cell type specific gene modules (calculated as the average depth-normalized read density of cluster specific gene markers (Table S1) followed by z-score scaling) as indicated for each cell type. GMM based cell annotations are indicated by colors on top of each heatmap that match the legend for cell types at the bottom of the figure. Multi- labelled cells based on GMM classification (FDR<0.05) are marked (*) and considered hybrids. (I-K) UMAP visualizations of GMM based cell state transitions (lines) transitioning between paired DCM (square) and RNA (circle) profiles for individual cells indicating their positions in UMAP space.

However, similar to the combinatorial indexing approach, we also observed several cells with DCM and RNA labels assigned to distinct major lineages that were not linked to mixed states. These occurred between secretory cell types and the absorptive lineage including cells with goblet- (n=2), Paneth- (n=10), tuft- (n=2), and EEC-like (n=6) DCM histories and current enterocyte RNA states (Fig. 2I, S5E, GMM FDR<0.05). Nevertheless, these were rare accounting for ∼7.8% of annotated histories in all enterocytes, and similar to the combinatorial indexing, most of these including all Paneth cells, were integrated in less mature compartments in UMAP space. Furthermore, both tuft cells also had histories consistent with cell cycle suggesting these could represent an earlier observed reserve stem cell population^36^ that consisted of rare cycling tuft cells^37^ .

In addition, we also noticed transitions occurring between most secretory cell types including goblet to Paneth and Paneth to goblet (Fig.2 F-K Fig. S5E, F; GMM FDR<0.05). The latter is compatible with previous lineage-tracing studies demonstrating that a small percentage of Paneth cells do not directly arise from a quiescent LGR5+ state^10,38^ as well as recent data showing that Paneth and goblet cells are district spatial manifestations of the same base cell that can transition based on spatial location^11^. Several rare discordant cells, including those spanning secretory and absorptive assignments, were robust to both GMM and data integration including Goblet to EEC (n=1), tuft to Paneth (n=1), and goblet to enterocyte (n=1) (Fig. 2I-K) representing clear switches in identity assignments.

### Intestinal differentiation has bidirectional properties

Besides inter lineage state discordance, we also observed immature TA and ISCs with prior secretory- or absorptive-lineage DCM histories including enterocytes (n=3), Paneth (n=7), goblet (n=3), tuft (n=5) and EEC (n=8) cells that were all marker gene verified (Fig. 3A, B; Fig. S5B; FDR<0.05). Several of these retained high-confidence assignments by both marker-gene scoring and data integration including enterocyte (n=2), goblet (n=1), tuft (n=2) and EEC (n=1) reversals (Fig. 3A, B; FDR <0.05). These cells are thus different from lineage primed cells which are defined as cells which retain a strong stem cell identity while expressing adult markers from multiple lineages and as such emerge as mixed states (further analyzed below). While the overall percentages of reversals were low (<5% of analyzed mature states), our data suggests that, under physiological conditions, intestinal epithelial cells can display bidirectional movements through differentiation state space.

**Fig. 3.**
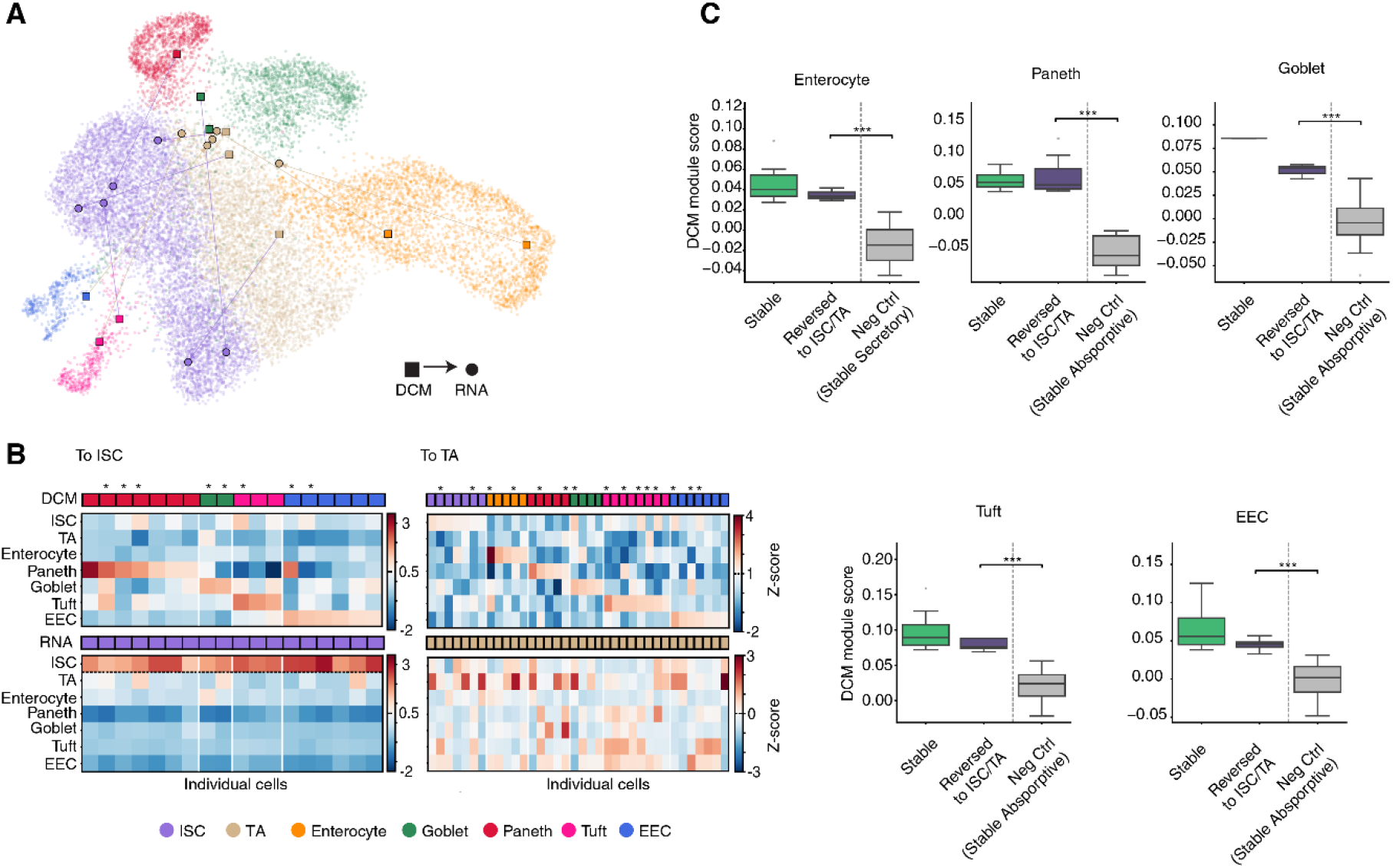
Modeling bi directional fate trajectories of differentiated cells in the mouse intestine. (A) UMAP visualizations of GMM based cell state transitions (lines) transitioning between paired DCM (square) and RNA (circle) profiles for individual cells indicating their positions in UMAP space. Colors match the legend for cell types at the bottom of the figure. (B) GMM based transitions. pDCM (top panel) and pRNA (bottom panel) data scaled across cell type specific gene modules (calculated as the average depth-normalized read density of cluster specific gene markers (Table S1) followed by z-score scaling) as indicated for each cell type. Each column is an individual cell. Multi-labelled cells based on GMM classification (FDR<0.05) are marked (*) and considered hybrids. GMM based cell annotations are indicated by colors that match the legend for cell types at the bottom of the figure. (C) DCM module scores of specific cell types grouped between stable cells, non-stable cells (cells reverting to ISC or TA) and negative control (stable absorptive cells (for Secretory scores) or stable secretory cells (for absorptive scores). Cells with mixed states are excluded. Each plot shows the module score of the corresponding gene set as indicated (*** (p < 0.001); * (p < 0.05), t-test).

To further characterize the nature of bidirectional lineage transitions, we compared lineage module scores between cell types that reversed differentiation direction to those that maintained a stable lineage identity. We found that module scores for cells that reversed their differentiation trajectory tended to be slightly lower compared to cells that stabilized their identity (Fig.3C). This suggests that cells that reverse differentiation may have failed to fully complete their lineage trajectory which is supported by many of these cells integrating in less mature compartments. Thus, rather than formal dedifferentiation, reversed trajectories may occur as a result of failures to complete or stabilize the differentiation process despite the activation of typical marker genes.

To analyze bidirectional dynamics in a less mature state, we next characterized mixed states containing TA or stem cell identities as part of their assignments (Figure 4A, B, n=65). These cells may be lineage primed states containing multiple signatures while including an immature identity^10,11^ or simply reflect intermediate state transitions. Of these cells, 22 cells completed differentiation consistent with their mixed profile, while 27 reverted to a non-mixed TA or stem cell state. No obvious differences in read densities were observed between the two populations (Figure 4C). This suggests that bidirectional movements become more prevalent in immature / primed states and drop in frequency as cells mature.

**Figure 4.**
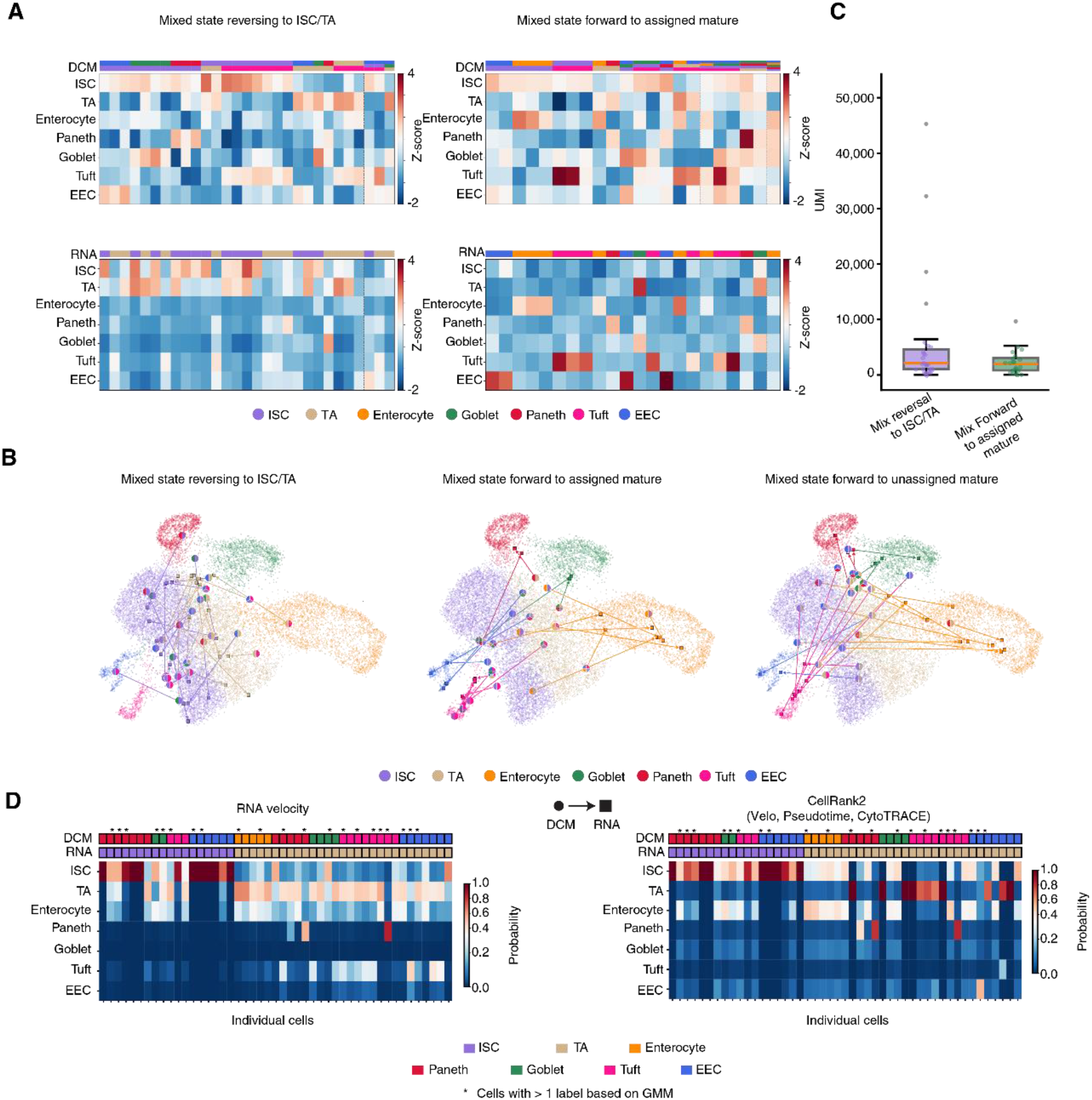
Bidirectional transitions in immature cells and future fate projections. **(A)** State transitions between cells with multiple GMM based state assignments (mixed states) that resolve to an ISC or TA state (left heatmap), or differentiate forward into an assigned mature state (right heatmap). pDCM (top panel) and pRNA (bottom panel) data scaled across cell type specific gene modules (calculated as the average depth-normalized read density of cluster specific gene markers (Table S1) followed by z-score scaling). Each column is an individual cell. GMM-based cell annotations are indicated by colors matching the cell-type legend at the bottom of the figure. Mixed states are shown with more than one color, based on assignment at Bayesian FDR < 0.05. (B) UMAP visualizing the reference data in background dots colored by their corresponding cluster, and on top highlighting the transitions of cells with mix states that resolve to ISC/TA (left UMAP); differentiate forward according to their mixed profile (middle UMAP); or cells that resolve towards an alternative state that was not part of the mixed state (right UMAP). DCM based assignments are circles and RNA assignments are squared. (C) Boxplot visualizing UMI counts for mixed cells resolving to immature (purple) or mature (green) states. The boxplot displays outliers (gray dots), the median (orange line), the interquartile range (box limits), and the whisker represent 1.5x of the interquartile range. (D) Heatmaps displaying the transition probabilities of individual cells displayed in Figure 3B, for transitioning to each possible macrostate (cell type). Measured transitions (DCM to RNA) for each cell are indicated by colors above each heatmap. Left heatmap shows transition probabilities based on RNA velocity analysis while the right heatmap shows probabilities by combining RNA velocity (30%), CytoTRACE (30%) and pseudotime (30%) modalities and cell connectivity (10%). Multi-labelled (mixed) cells based on GMM classification (FDR<0.05) are marked (*).

### Future-state projections support a model for lineage exploration

As bidirectional movements during differentiation provide a simple explanation for the apparent lineage switches that we observed between adult states, we asked whether current ISC/TA-like cells carrying DCM histories consistent with a mature state were predicted to re-enter their original trajectory or adopt alternative lineages. We first used RNA velocity analysis which predicts short term state transitions by comparing spliced and un-spliced RNA^39,40^. Cluster level predictions were analyzed to forecast state transitions on a single cell basis (Fig. S6, Fig. 4D). This revealed that most cells (95%) were projected to either stay immature or revisit the differentiation path from which they came. However, several cells with secretory histories that were in an immature state (ISC/TA) were now projected to transition towards the enterocyte lineage through a TA state and others were predicted to change secretory lineage. To confirm these observations across a longer projected timeframe we used CellRank2, a model that integrates three analyses, CytoTRACE ^41^, pseudotime ^42^ and RNA velocity analysis ^39,40^, to estimate fate probabilities (Fig. S6, Fig. 3D) ^43^. We predicted fate outcomes at cluster level for each of the three analyses separately showing the main directionality of cluster transitions (fig. S6), and then merged the three modalities in a combined analysis (fig. S6). These analyses largely confirmed the RNA velocity data with several secretory reversals now being projected towards the enterocyte lineage.

For cells originating from TA cells, we observed similar probabilities of differentiation occurring towards either ISCs or enterocytes in support of differentiation trajectories being bidirectional (Fig. 4D). This is also consistent with previous studies demonstrating that some cells in the TA compartment can migrate back to the crypt and act as primary progenitors^44^.

To move beyond predictions, we next analyzed lineage primed or mixed intermediate cells most of which resolved towards their primed state or reverted back (Fig. 4A,B). However, 16 of these changed directions in a way that was inconsistent with their lineage primed history. Again, these included only secretory to secretory (n=7), or secretory to absorptive switches (n=9).

These analyses suggest a model in which cells that fail to stabilize their initial lineage program may in rare cases be able to resolve toward lineages distinct from their initial DCM-inferred lineage-associated program, providing a simple explanation for the observed inter lineage state discordance (Fig 2F-K).

### Early neural differentiation shows similar bidirectional state histories

To study the dynamics of cell state changes in a second system that was even less mature than intestinal mixed states, we generated neural stem cells from mESCs containing the DCM- Polr2b system, regulated by a dTAG system to enable its prompt destruction (Fig. S7)^45^. We used these to track transcriptome histories during early neural differentiation in vitro ^45,46^. Differentiation into immature neurons (INs) and immature glial cells (IGs) was benchmarked across 6 days following EGF removal using scRNAseq and multiome analysis (Fig. S8). An immature neural cell type was present from day two onward that had lost the *Hes5* radial glial marker and obtained immature neuronal markers Gria2 and Class III β-tubulin (*Tubb3*) ^46–48^. INs were also positive for doublecortin (*Dcx*) and dorsal marker *Sox6* ^49^. As expected, a separate cluster expressed *Prrx1* and *Twist1* resembling immature glial cells (IG), was also found^46,50^. To trace cell state transitions in this system, we exposed cells to doxycycline for 16 hours to induce transgene expression at day 2 followed by dTAG mediated destruction of the fusion protein. These cells were then chased for 24 hours and analyzed using scMeD&RNA-Seq (Fig. S9A). We detected an average of 3,150 genes by mRNA, 1,971 genes with methylated DCM sites and up to 87,596 methylated CpGs per cell depending on sequencing coverage (Fig. S9C-F). Following BBKNN integration, DCM based cell identities were determined using GMM (FDR<0.05) using cell type specific marker gene sets (Fig.5A, B; Fig S9G, H)

**Fig. 5.**
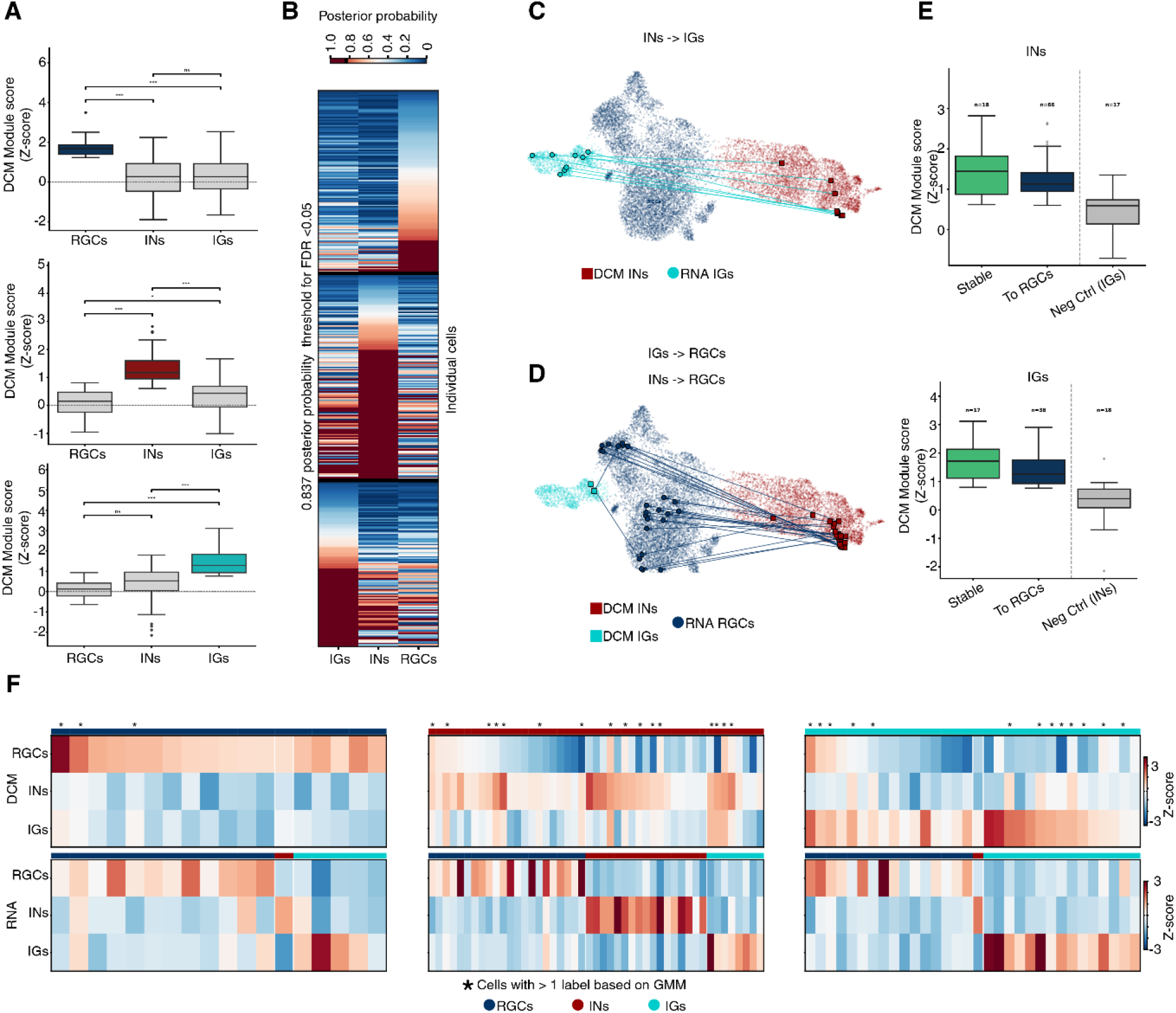
Bidirectional properties of early neural differentiation. (A) DCM module Z-scores for marker gene sets specific to each cluster (cell type) calculated for all indicated cell types. Each panel indicates a separate cell type specific gene module set. Z-scores indicate how strongly the cell’s gene expression pattern deviates from the average expression pattern across all cells in the dataset. The colored boxes highlight the cell type upon which module genes were selected matching the colors in C and D. t-tests are FDR adjusted (*** (p < 0.001); ** (p < 0.01); * (p < 0.05); ns (p > 0.05)). (B) Heatmap showing module score based posterior probabilities generated using Gaussian Mix Modelling (GMM, >0.85 posterior probability). (C) UMAP showing the transitions from filtered INs to IGs and vice versa. Starting states (DCM) are indicated by squares while end states (RNA) are represented by circles. (D) UMAP as in C but showing cells reverting from either INs or IGs back to more immature RGCs. Cells are colored by their respective cell types. (E) DCM module scores of specific cell types grouped between stable cells, non-stable cells (cells reverting to RGCs) and negative control (stable INs for IGs and vice versa). Each plot shows the module score of the corresponding gene set as indicated. (F) Z-score DCM and RNA module scores are calculated as the average depth-normalized read density of cluster specific gene markers (RGCs, n=399; IN, 567; and IG, n=1,089) for both pRNA and pDCM data, following by z-score scaling across gene modules (gene sets representing a cluster or cell type) within each cell to highlight the relative contribution of each gene module. Top and bottom panel show DCM and RNA profile scores as indicated. The heatmaps focuses on filtered cells transitioning between RGCs (left panel), INs (middle panel) and IGs (right panel) other lineages. Multi-labelled cells based on GMM classification (FDR<0.05) are marked (*).

Consistent with our results from the mouse intestine, we observed rare cells with IN-like DCM histories and IG-like current RNA states (n=9; Fig. 5C and F). In addition, we also observed bidirectional transitions, with immature radial glial like cells carrying IN- or IG-like histories (n=38; Fig. 5D and F). Cells that reversed their lineage trajectory were found to have slightly reduced marker gene activation compared to cells that remained stable (Fig. 5E, Fig S9I-J). This suggests that incomplete lineage stabilization and bidirectional lineage movements are not restricted to the intestinal epithelium.

## Discussion

In the intestine, the reliance on lineage tracing as a single tool to rebuild lineage trajectories has yielded conflicting models on how this tissue is organized^9,29^. By reconstructing genome-wide prior transcriptional-state signatures rather than relying on individual marker activity, our approach now provides a broader layer of temporal information. This analysis suggests that during differentiation some cells may transiently switch between lineage-associated programs without fully stabilizing these transitions, resulting in apparent reversals through state space. This is different from forward lineage priming in which cells retain a stem cell identity while expressing markers of multiple differentiated cells^34^. In our data these lineage primed cells show up as mixed states with both stem and adult signatures. We find that these mixed states similarly transition in a bidirectional fashion rather than resolving into one chosen state. However, cells that reverse through state space form a near mature state are rare within the population. Nevertheless, given the total number of cells in the intestine, thousands of cells are likely to make such discordant movements every day. Such cells may subsequently re-enter the same trajectory or, in rare cases, resolve toward an alternative lineage state. This would explain some of the similarly rare inter lineage transcriptional state switches between lineages that we observed.

While orthogonal approaches covering two time points in single cells are not yet available save for classical lineage tracing studies, these are consistent with our analysis as rare ribbons that indicate expression of canonical lineage markers in bona fide stem cells are observed in most of the lineage specific CRE-markers used including those to identify secretory progenitors^33,51,52^. Our data suggests that at least some of these may reflect rare physiological transitions that take cells back to an ISC state rather than spurious CRE activity.

Whether such bidirectional movements during differentiation reflect inherently noisy stochastic dynamics or specific biological mechanisms, such as retrograde movements^53^, morphogen pulses^54^, transient cell-cell contacts, or a combination of these, remains unclear. The transient nature of any of these signals may explain why cells vary in their ability to fully execute a differentiation step. It is also unclear to what extend fluctuations in transcriptomes translate directly into permanent changes at the cellular level as proteins are typically longer lived than mRNA. As such establishing a new proteome that overcomes a previous lineage may entail several back-and-forth switches (i.e. differentiation attempts) at the transcriptomic level. Our observation of similar bidirectional state histories in a model of early neural differentiation suggests that the properties observed in differentiating epithelial tissues may be shared during early developmental processes. Nevertheless, the generalizability of these concepts needs further exploration. If such bidirectional movements prove to be widespread, differentiation may be better viewed not simply as a downhill trajectory but rather as an uphill process in which cells must stabilize identity against competing or incompletely resolved lineage programs. Future work is needed to firmly establish these concepts across development as well as the potential role these processes may play in lineage infidelities and the emergence of cancer^8^ .

## Methods

### Single cell state tracing in the mouse intestine

#### DCM-Polr2b transgenic mice

Animals were bred and maintained in the Erasmus MC animal facility (EDC) under conventional specific pathogen-free conditions. DCM–Polr2b:m2rtTA mice were described previously ^25^. Transgene expression in DCM–Polr2b:m2rtTA mice was induced by the addition of doxycycline (dox; Sigma-Aldrich, D9891) to the drinking water of mice (2 mg/ml, 2% sucrose (Sigma-Aldrich, 84097)) for 48h. Mice were sacrificed using cervical dislocation. All animal experiments were approved by the Dutch Central Committee on the Ethics of Animal Experiments (AVD10100202115681).

#### Isolation of cell populations from the small intestine and FACS sorting

Small intestine was dissected from sacrificed mice. The isolation of cell suspensions from the jejunum for enterocytes was performed as described previously ^25^. Briefly, the jejunum was isolated from the small intestine, the adjacent fat tissue was removed and the tissue was washed with PBS while shaking on ice. The jejunum was opened using surgical scissors, cut into small pieces (∼1cm long) and incubate on ice for 30 minutes in 2mM EDTA/PBS. After two PBS washes, the tissue pieces were pipetted vigorously 12 times in a 10mL pipette tip. The supernatant, containing villi, was incubated at 37°C for 30 minutes while shaking at 120 cycles/min in 1x Dispase II (Gibco, 17105041) to obtain a single cell suspension. The cells were filtered with a 40 µm strainer and collected for staining in 2% FCS/PBS staining buffer. For enterocyte purification, cells were stained with conjugated (eFluor 450) monoclonal antibody for EpCAM (Thermofisher, 48-5791-82; 1:50) and Cy5 conjugated polyclonal antibody for GLUT-2 (Bioss, bs-0351R-Cy5; 1:20) for 30 minutes on ice. Double-positive cells were FACS sorted using a BD FACSAria II version 9.0.1 (BD Biosciences). A total of 500,000 cells were collected to proceed with the combinatorial indexing protocol (see below).

For other cell populations, crypt isolations were performed as described in previously^55^. Briefly, the jejunum was collected from sacrificed mice and surrounded fat tissue was removed. Tissue was cleaned with PBS and opened using scissors. The villi were scraped off with a glass slide and the intestine was cut in 5mm pieces. The pieces were washed with PBS 3-5x and incubated for 30 minutes at 4°C with 6mM EDTA in PBS. Subsequently, the pieces were transferred to PBS and homogenized by pipetting vigorously 10-15 times using a 10mL pipette to dissociate the crypts from the tissue. The supernatant was filtered through a 100µm filter. Tissue pieces were taken from the filter and the process was repeated until the PBS was clear. The collected supernatant was centrifuged at 300g for 5 minutes at 4°C and the pellet was resuspended in 1mL DMEM/F12 (12634010, ThermoFisher). DNaseI (10104159001, Merck) was added to a final concentration of 0.5mg/mL, and the sample was incubated for 10 minutes at room temperature (RT). The sample was resuspended gently with 9mL of DMEM/F12 and centrifuged at 80g for 5 minutes at 4°C. The pellet was resuspended in 1mL of pre-warmed TrypLE containing 0.5 mg/mL DNaseI, and incubated for 3 minutes at 37°C in a water bath while homogenizing with a P1000 pipette every minute. After adding 9mL DMEM/F12 samples were centrifuged at 300g and 4°C in a 15mL falcon. Pellets were resuspended in 5% FBS/PBS and stained with antibodies for CD31-BV421 (BD Biosciences, 563356), CD45- BV421 (BD Biosciences, 563890), and TER119-BV421 (BD Biosciences, 563998), each at a final concentration of 2µg/mL), for 30 minutes on ice. DAPI staining was performed at a final concentration of 1µg/mL to exclude dead cells. The cells were stained for CD24-APC (Sony Biotechnology, 1109070; final concentration: 2µg/mL), cKit-PE (BioLegend, 105808; final concentration: 2µg/mL) to sort for multiple populations, using BD FACSDiscover™ S8 Cell Sorter (BD Biosciences)^55,56^. For plate-based sequencing, cells were sorted in a 384-well plate and doublets were visualized using BD CellView Image Technology and removed from the plate before sequencing. For bulk analysis or combinatorial indexing cells were sorted in PBS. FACS analysis was visualized with BD FACSChorus^TM^ Software (v 5.4.0).

### Single-cell MeD-Seq using combinatorial indexing (sciMeD-Seq)

#### Intestine nuclei isolation, fixation and permeabilization

After sorting enterocytes using EpCAM and GLUT-2, the cells were spun down at 500g for 5 minutes and washed in 1mL PBS. The cells were counted with the manual counter giving a total of 540,000 cells. Nuclei were isolated by incubating 500µL hypotonic lysis buffer together with 0.025% Igepal on ice for 5 minutes. The nuclei were spun down (500g, 5min, at 4°C) and gently resuspended in 500µL hypotonic lysis buffer containing a final concentration of 0.752% using 16% formaldehyde (Merck, F1635). The nuclei were incubated at room temperature for 10 minutes, followed by the addition of an equal volume of 2.5M glycine (Merck, G8790) relative to the added formaldehyde (23.5µL), to quench the fixation process, and incubated for 5 minutes on ice. Nucleosomes were depleted by resuspending the pellet in 500µL 1x 2.1 NEB buffer (NEB, B7202S) containing 0.3% SDS (Bio-Rad, 1610418) and incubating for 20 minutes at 37°C. The nuclei were spun down and washed with 500µL 0.3M SPBSTM^57^ (11.4% sucrose, 0.1% of Triton-X100, 1x PBS, 3mM MgCl_2_) and a second time with 500µL 1x rCutSmart buffer (rCS; NEB, R0663S) in 0.3M SPBSTM. Intestinal nuclei were further processed using LpnPI digestion described below (section: *LpnPI digestion of methylated DNA and cleanup)*.

#### ESC nuclei isolation, fixation and permeabilization

For cultured cells, single cell ESC suspensions were collected following trypsinization and washing with PBS. Cells were counted using a TC20 automated cell counter (Bio-rad) and 5 million mESCs were collected and spun down (500g, 5min at 4°C). Nuclei were isolated by incubating in 5mL hypotonic lysis buffer^57^ (1x PBS, 11.4% sucrose (Merck, S8501) and 3mM MgCl_2_ (Merck, 68475) in water) together with 0.025% Igepal (Sigma, I8896), on ice for 10 minutes. Nuclei were spun down (500g, 5min, at 4°C) and gently resuspended in 1mL 0.3M SPBSTM (11.4% sucrose, 0.1% of Triton-X100, 1x PBS, 3mM MgCl_2_) containing 12.5mM lithium diiodosalicylate (Sigma, D3635) for nucleosome depletion. Nuclei were then incubated on ice for 5 minutes, followed by centrifugation for 5 minutes at 500g and 4°C. For each sample, 100µL 50mg/mL DSP in 4mL ice cold methanol was gradually mixed with the pelleted nuclei and fixation on ice was done for 15 minutes on a gently rocking surface. Two equal volumes of 0.3M SPBSTM was added gradually to each reaction and the nuclei were spun down for 5 minutes at 500g. The supernatant was removed, and nuclei were washed in 1mL 0.3M SPBSTM and a second time in 800µL 1xrCutSmart buffer (rCS; NEB, R0663S) in 0.3M SPBSTM. Similar to intestinal cells, ESC nuclei were further processed using LpnPI digestion described below.

#### LpnPI digestion of methylated DNA and cleanup

Nuclei were counted using the TC20^TM^ Automated cell counter (Bio-Rad) and resuspended in 1x rCS buffer containing 0.3U/µL of LpnPI aiming for a final concentration of 2,500 nuclei/µL in a total reaction volume of 50µL. The reaction was incubated at 37°C overnight. The nuclei were collected, resuspended in 200µL 0.3M SPBSTM, and counted for use in combinatorial adapter ligation reactions.

#### Adapter ligation

Nuclei were resuspended at a final concentration of 2,500 nuclei/10µL ligation buffer (1x NEB T4 ligase buffer (NEB, B0202S) prepared in 0.3M SPBSTM), and split equally in an 8- well PCR strip (BIOplastics BV, B57801). The nuclei were divided into the 1st 96 well plate (10µL/well; Thermofisher, 165305). Subsequently, 1.5µL of T4 mix buffer (ligation buffer containing 0.45U/µL T4 ligaseand 1x NEB T4 buffer) and 0.5µL of 100µM ‘Odd’ adapters (Table S2. sciMeDseq adapters, IDT) were added. The plate was sealed and incubated at room temperature for 30 minutes followed by 5 minutes on ice. The nuclei were pooled and washed 2x with 1mL 0.3M SPBSTM and spun down at 500g for 5min at 4°C. After the last wash, the supernatant was removed and previous steps were repeated with the second and third round of adapters (Table S2. sciMeDseq adapters, IDT). After the last round of ligation, the cell pellet was resuspended in 20µL 0.3M SPBSTM and the nuclei were counted. The nuclei were separated in pools of 500-1000 nuclei in 6µL 0.3M SPBSTM for protease treatment with 2µL 1.07AU/mL protease (Qiagen, 19155) at 55°C for 1h. Following heat inactivation at 75°C for 30 minutes, lysates were stored at −80°C until further processing.

#### Linear amplification

The digested nuclei were mixed with 0.75µL of 50µM P5 partial NEBNext index with random nanomers (Table S2. sciMeDseq libraryprep oligos, IDT) and incubated in the following program: 95°C for 3 minutes; gradually cool down (0.1°C/s) to 20°C. Subsequently, 1.6µL gap extension master mix (1x NEBuffer 2, 0.5mM dNTP, and 2U/ µL Klenow Fragment Exo (3’→5’ exo-); NEB, M0212L)) was added and incubated at 37°C for 10 minutes followed by incubation at 75°C for 10 minutes. After cooling on ice, in vitro transcription was performed following Hi Scribe T7 Quick RNA synthesis Kit (NEB, E2040S). Briefly, 12µL IVT master mix (2µL T7 Polymerase, 10µL NTP buffer mix) was added followed by an overnight incubation step at 37°C. 1µL of 2U/µL DNaseI (NEB, E2040S) in 29µL H_2_O was added to remove original DNA template by incubating for 15 minutes more at 37°C. Amplified RNA (aRNA) was then purified with 0.7x SPRI beads (Beckman Coulter, B23317) following the manufacturer’s instructions. The beads were then eluted in 11µL H_2_O to continue with the Reverse transcription step.

#### Reverse Transcription

The reverse transcription was performed following the SuperScript^TM^ IV Reverse Transcriptase (RT) kit (Thermofisher, 18090010) with a few modifications. Eluted aRNA was mixed with 1µL of 50µM partial P5 index primer (Table S2. sciMeD-Seq libraryprep oligos) and 1µL of 10mM dNTPs. The reaction was incubated at 70°C for 5 minutes and at 90°C for 20 seconds followed by cold shock on ice. 7µL of RT master mix (1x SSIV buffer, 5mM DTT, 1U RNaseOUT (Invitrogen, 10777019) and 10U SSIV adjusted to the 20 µL final volume) was added to the sample. The RT reaction was incubated in a thermocycler using program: 55°C for 15 minutes, 60°C for 10 minutes, 65°C for 12 minutes, 70°C for 8 minutes, 75°C for 5 minutes and 80°C for 10 minutes. To remove the aRNA template, 0.5µL of Rnase H (NEB, M0297) and 0.5µL RNase A (Invitrogen, AM2270) were added and samples were incubated at 37°C for 30 minutes.

#### PCR amplification and Sequencing

To avoid under or over amplification of the sample, we used 1/10^th^ of the sample for qPCR to obtain an estimate for the number of PCR cycles. GoTaq® qPCR Master Mix kit (Promega, A6002) was used according to the manufacturer instructions using P5 and P7 NEBNext Multiplex Oligos for Illumina (E7600S, NEB, Table S2. Illumina index primers). These primers anneal to the adapter barcode and to the incorporated partial P5 primer sequence. The samples were amplified in the thermocycler: 98°C for 30s; 40 cycles of (98°C for 10s, 64°C for 30s, 72°C for 30s); 72°C for 1 min. PCR cycle numbers were determined by using the following formula: *Estimated PCR cycles* = *cq value* − 3, and cycle number estimates were used below.

Half of the RT reaction was used for PCR amplification, combining NEBNext High-Fidelity PCR Master Mix (NEB, M0541S), and P5 and P7 indexes in a final concentration of 0.5 µM each, using the following cycles: 98°C 30s, an estimated cycles of (98°C 10s, 65°C 30 s, 72°C 30s), 72°C 1 minute and 4°C on hold. dsDNA was purified with 0.75x SPRI, as previously described according to manufacturer instructions. The library was diluted in 12µL nuclease-free water and fragment size and concentration was determined using a bioanalyzer. Fragments below 200bp were removed using 0.7x AMpureXP beads (Beckman Coulter, A63881) to avoid adapter dimer sequencing. The library was sequenced using paired-end (PE) sequencing at either 150bp or 100bp using the NOVAseq6000 platform with 10-35% Phix genome added to increase complexity of the library. Dual indexed samples were demultiplexed using bcl2fastq (Illumina).

### Combined MeD-Seq & RNA-Seq in single cells using plate-based reaction miniaturization

#### Single cell sorting into plates

Single live cells were sorted into low bind 384-well plates (Eppendorf, 0030129547) using the BD FACSAria II Cell sorter System or FACSDiscover (BD Biosciences), with plate cooling as described above in the section: *Isolation of cell populations from the small intestine and FACS sorting.* A dilution of 1:1000 zombie R718 (Biolegend, 423116) or 10µg/mL DAPI (D3571, ThermoFisher) was used to label dead cells. Before sorting, the plates were prepared by adding 1µL mild lysis buffer (0.1% Igepal (Sigma, I8896), 1U/µL Recombinant ribonuclease inhibitor (Takara, 2313B)) and 2µL RTL plus buffer (Qiagen, 1053393) using multichannel pipettes. Single cells were sorted in each well and after sorting, the plate was spun down 1,000g for 1 minute at 4°C. Plates were stored at −80°C for later processing. Single Cell content for each well was verified using BD CellView Image Technology from FACSDiscover.

#### Separation of mRNA and gDNA

To capture mRNA from cell lysates a strategy was designed based on scNMT-seq^58^. For a full 384-well plate, 225µL Dynabeads™ MyOne™ Streptavidin C1 (ThermoFisher, 65001) was transferred to a 1.5 mL tube and placed on a magnetic rack (Invitrogen). The supernatant was removed, and beads were washed with 400µL solution A (0.1M NaOH, 0.05M NaCl in nuclease-free water). The previous step was repeated one more time, followed by a wash with 200µL solution B (0.1M NaCl in nuclease-free water). The beads were resuspended in 225µL 2x B&W buffer (0.01M Tris-HCl pH 7.5, 1mM EDTA, 2M NaCl in nuclease-free water) and 225µL 100µM oligo-dT-SS3 (Table S3. scRNA-MeDseq & library oligos, IDT) was added. The bead-oligo-dT mix was incubated by gentle rotation for 15 minutes at room temperature on a roller in a 50mL falcon tube. Conjugated beads were washed three times with 400µL 1x B&W buffer and resuspended in bead resuspension buffer (1x RT buffer, 1U/µL RNAse inhibitor (RRI)). Beads (4µL per well) were added to the 384-well plate with the lysed cells and resuspended manually using Rainin automatic pipette 0.5-10 µL LTS (Mettler Toledo). The plate was incubated at room temperature for 10 minutes followed by a centrifugation step of 1,000g for 1 min at room temperature. The plate was loaded on a plate magnet (SPT Labtech, 3268-02008) for 2 minutes and the supernatant containing the gDNA was transferred to a new plate manually (gDNA plate) using a multichannel pipette. The beads were washed with 5µL G&T wash buffer (0.05M Tris-HCl pH 8.3, 0.075M KCl, 3mM MgCl_2_, 0.01M DTT, 0.5% Tween-20 in nuclease-free water), followed by an incubation of 5 minutes at room temperature. After a centrifugation step followed by magnetic attachment of the beads, the supernatant was also transferred to the gDNA plate. With a total of 12µL, the gDNA plate was stored at −20°C for subsequent processing using plate-based scMeD-seq. The plate with the beads-conjugated mRNA was processed using scRNA-seq based on SMART-Seq3.

#### scRNA-Seq (SMART-seq3 based), Reverse Transcription and PCR

Reverse transcription reaction was performed by resuspending the beads in 2µL RT master mix (1mM dNTPs, 25mM Tris-HCl pH8.3 (Bioworld Technology, 1185-53-1), 30mM NaCl, 2.5mM MgCl_2_, 1mM GTP (ThermoFisher, R0461), 2µM TSO (Table S3. scRNA- MeDseq library oligos), 0.5U/µL RRI, 5% PEG 8000 (Molecular dimensions, P1458), 8mM DTT (Invitrogen, Y00147), 2U/µL Maxima RT (ThermoFisher, EP0752)) followed by a brief centrifugation step (100g for 15s). The plate was incubated in a thermocycler (Bio-Rad) using the following program: 42°C for 90 min; {50°C for 2min followed by 42°C for 2min;10 cycles}; 85°C for 5min; hold at 4°C. After the RT reaction, the plate was spun down at 1,000g for 1 minute at room temperature. The beads with the RT reaction were resuspended in 3µL PCR mastermix (1x HiFi with Mg. KAPA High Fidelity buffer (Roche, 7958897001), 0.5mM MgCl2, 0.5mM dNTPs, 0.5µM Fwd PCR primer, 0.1µM Rev PCR primer (Table S3. scRNA- MeDseq library oligos) and 0.02U/µL HiFi HotStart KAPA polymerase (Roche, 7958927001)) followed by amplification in a thermocycler (Bio-Rad): 98°C for 3 min; {98°C for 20s, 65°C for 30s; 72°C for 4min; 28-30 cycles}; 72°C for 5min; 4°C hold.

#### scRNA-Seq (SMART-seq3 based), Ampure XP beads purification

The AMpure XP beads (Beckman Coulter, A63881) were used according to manufacturer instructions and minor modifications were made. Briefly, a bead to sample ratio of 0.7x was used followed by an incubation of 10 minutes at room temperature. The plate was placed in the 384 PCR Plate Magnetic Bead Clean-Up Block (SPT Labtech, 3268-02008) for 2 minutes to attach the beads to the magnet. The supernatant was then removed and beads were washed with 80% ethanol. After removing the ethanol, the beads were dried for 2 minutes, resuspended and incubated for 5 minutes in 4µL nuclease-free water. The beads were then attached to the magnetic block and the eluted sample was transferred to a new plate (cDNA plate). The fragment distribution and concentration of eleven random wells were used as input in the bioanalyzer (Agilent), using the High Sensitivity DNA kit (Agilent, 5067-4626). The average concentration of the 11 wells was used as a reference to dilute the full plate into roughly 750pg/µL of cDNA which was transferred to a new plate (normalized plate). The normalized plate was generated only when the average concentration exceeded 750pg/µL. This plate was either stored at −20°C or processed for subsequent tagmentation.

#### scRNA-Seq (SMART-seq3 based), Tagmentation and sequencing

The Mosquito LV liquid handler was used to dispense 300pg cDNA from the normalized cDNA plate into a new plate (tagmentation plate), either from the normalized cDNA plate or, for non-normalized samples, directly from the original plate based on the measured cDNA concentration. The tagmentation plate was pre-processed to contain 1.2µL tagmentation master mix (0.8µL 2x Tagment DNA buffer, 0.1µL Amplicon Tagment Mix, 0.3µL H_2_O), from the Nextera XT DNA Library Preparation Kit (Illumina, FC-131-1024). The plate was briefly spun down and incubated for 10 minutes at 55°C. Subsequently, 0.4µL of 0.2% NT buffer (Illumina, FC-131-1024) was dispensed manually into the tagmentation plate followed by incubation for 5 minutes at room temperature to stop the reaction. The index primers (Table S3. scSMARTseq3 index primers, IDT) were dispensed using the Mosquito to a final concentration of 0.1µM, followed by 1.2µL of Nextera PCR Mastermix (Illumina, FC-131-1024) or 2.4µL KAPA HiFi HotStart ReadyMix (Roche, 9420398001) by manual dispensing. Amplification was performed using the following programs: for Nextera PCR Master Mix: 72 °C for 3 min; 98 °C for 3 min; 16 cycles of 98 °C for 10 s, 55 °C for 30 s, and 72 °C for 30 s; final extension at 72 °C for 5 min; hold at 4 °C. For KAPA HiFi HotStart ReadyMix: 72 °C for 3 min; 98 °C for 3 min; 16 cycles of 98 °C for 20 s, 63 °C for 15 s, and 72 °C for 30 s; final extension at 72 °C for 5 min; hold at 4 °C. Tagmented cDNA was purified using 0.7x AmpureXP beads as described above. After analyzing tagmented profiles using the bioanalyzer, samples were pooled manually and purified again with 0.7x AMpure XP beads and washed once with 80% ethanol. The beads were diluted in 35µL, and after separation from the sample (supernatant) using a magnetic rack, the sample was checked in the bioanalyzer and sent for sequencing. The cDNA was sequenced with 10-35% Phix genome to increase complexity of the library in Novaseq 6000 (Illumina) obtaining 100 or 150bp paired end reads. At least a depth of 50,000 reads per cell was achieved to obtain enough information per cell.

#### scMeD-seq: LpnP1 digestion and adapter ligation

gDNA plates were thawed on ice and purified with 0.7x Ampure XP beads as described above. Beads were resuspended in 1.2µL nuclease-free water to elute DNA. Using the Mosquito liquid handler, gDNA containing supernatant was transferred to a new plate. The I-DOT liquid handler (Dispendix) was used to dispense the master mixes of each of the reaction’s components below while the Mosquito LV liquid handler was used to dispense the adapter barcodes from stock plates (Table S3. scMeD-Seq adapters). Plates were spun down for 1 min at 1000g after each master mix dispensing step, and cooled on ice for 2 minutes, following each incubation. Methylated gDNA was digested by dispensing 300nL of LpnPI digestion mix into each well (1x rCS buffer, 0.073U/µL LpnPI, 0.5µM Enzyme activator solution). The reaction was stopped by incubating the plate at 65°C for 20 minutes. After digestion, the adapters (Table S3. scMeD-Seq adapters) were dispensed at a final concentration of 10nM. Adapters contained a T7 promoter for in vitro linear amplification, a 3 nucleotide UMI to obtain the number of captured unique molecules, and an 8-nucleotide barcode for cell identification. 40nL adapter dispension was followed by adding 400nL T4 ligation master mix (1x NEB ligase buffer, 0.06U/µL T4 ligase (ThermoFisher, EL0012)). The plate was incubated overnight at 16°C in a PCR machine (Bio-Rad) and either processed further or stored at −20°C.

#### scMeD-seq: Sample pooling and cleanup

The gDNA plate was turned over and spun down to pool adapter ligated gDNA in a reservoir. Pooled sample was recovered in 1.5mL Lobind tubes, centrifuged, and purified using 8x diluted AMpureXP beads in bead buffer (20% (wt/vol) PEG 8000, 10 mM Tris-HCl, 1 mM EDTA, 2.5M NaCl and 0.05% (vol/vol) Tween-20) at a 0.7x bead-to-sample ratio, following the manufacturer’s protocol. The samples were eluted in 6µL nuclease-free water for linear amplification.

#### scMeD-seq: Linear amplification

The eluted samples were mixed with 50µM partial P7 NEBNext primer with random nanomers (Table S3. scRNA-MeDseq library oligos) and incubated in the following program: 95°C for 3 minutes; gradually cool down (0.1°C/s) to 20°C. Subsequently, 1.6µL gap extension master mix (1x NEBuffer 2, 0.5mM dNTP, and 2U/ µL Klenow Fragment Exo (3’→5’ exo-); NEB, M0212L)) was added and incubated at 37°C for 10 minutes followed by incubation at 75°C for 10 minutes. After cooling on ice, in vitro transcription was performed following Hi Scribe T7 Quick RNA synthesis Kit (NEB, E2040S). Briefly, 12µL IVT master mix (2µL T7 Polymerase, 10µL NTP buffer mix) was added followed by an overnight incubation step at 37°C. 1µL of 2U/µL DNaseI (NEB, E2040S) in 29µL H_2_O was added to remove original DNA template by incubating for 15 minutes more at 37°C. Amplified RNA (aRNA) was then purified with 0.7x SPRI beads (Beckman Coulter, B23317) following the manufacturer’s instructions. The beads were then eluted in 11µL H^2^O to continue with the Reverse transcription step.

#### scMeD-seq: Reverse Transcription

The reverse transcription was performed following the SuperScript^TM^ IV Reverse Transcriptase (RT) kit (Thermofisher, 18090010) with a few modifications. Eluted aRNA was mixed with 1µL of 50µM 1^st^-PCR-P7 primer and 1µL of 10mM dNTPs. The reaction was incubated at 70°C for 5 minutes and at 90°C for 20 seconds followed by cold shock on ice. 7µL of RT master mix (1x SSIV buffer, 5mM DTT, 1U RNaseOUT (Invitrogen, 10777019) and 10U SSIV adjusted to the 20 µL final volume) was added to the sample. The RT reaction was incubated in a thermocycler using program: 55°C for 15 minutes, 60°C for 10 minutes, 65°C for 12 minutes, 70°C for 8 minutes, 75°C for 5 minutes and 80°C for 10 minutes. To remove the aRNA template, 0.5µL of Rnase H (NEB, M0297) and 0.5µL RNase A (Invitrogen, AM2270) were added and samples were incubated at 37°C for 30 minutes.

#### scMeD-seq: qPCR QC, PCR amplification and Sequencing

To avoid under or over amplification of the sample, we used 1/10^th^ of the sample for qPCR to obtain an estimate for the number of PCR cycles. GoTaq® qPCR Master Mix kit (Promega, A6002) was used according to the manufacturer instructions using 0.5µM P5-1stPCR and 0.5µM P7-1stPCR (Table S3. scRNA-MeDseq library oligos, IDT). These primers anneal to the adapter barcode and to the incorporated partial P7 primer sequence. The samples were amplified in the thermocycler: 98°C for 30s; 40 cycles of (98°C for 10s, 64°C for 30s, 72°C for 30s); 72°C for 1 min. PCR cycle numbers were determined by using the following formula: *Estimated PCR cycles* = *cq value* − 3, and cycle number estimates were used below.

After determining the number of PCR cycles, a first PCR was performed to add Illumina adapters by using half of the reverse transcribed sample with 25µL 2x NEBNext High-Fidelity PCR Master Mix (NEB, M0541S), 2µL 10µM partial P5 and P7 primers (P5-1stPCR, P7- 1stPCR; Table S3. scRNA-MeDseq library oligos, IDT), and 10.5µL nuclease-free water. The thermocycler program applied was: 98°C for 30s; {98°C for 10s, 60°C for 30s; 72°C for 30s; *Estimated PCR cycles*/2}; 72°C for 1min; 4°C hold. After amplification, the pooled sample was purified with 0.75x AMPureXP beads. The pooled sample was eluted from the beads in 21 µL of nuclease-free water.

A second PCR was used to add the Illumina indexes for sequencing, using NEBNext® Multiplex Oligos for Illumina® (Dual Index Primers Set 1; NEB, E7600S). The pooled sample was mixed with 25µL 2x NEBNext High-Fidelity PCR Master Mix, 2µL 10 µM i5 index and 2µL 10µM i7 index. The pooled sample was amplified using: 98°C for 30s; {98°C for 10s, 65°C for 30s; 72°C for 30s; *Estimated PCR cycles*/2}; 72°C for 1 min; 4°C hold. Following amplification, the pooled sample was purified with 0.7x AMPureXP beads as described above. The beads were eluted in 12µL of nuclease-free water. The sample was run in a High Sensitivity chip using the bioanalyzer to measure the fragment distribution and concentration. Pooled samples were sequenced on the Illumina NovaSeq 6000 platform paired-end with a read length of 100bp or 150bp. For gDNA methylation and RNA samples were sequenced at a depth of between 0.5-1 million and 0.1-0.5 million reads per cell, respectively.

### Preprocessing combinatorial indexing-based sciMeD-Seq and plate-based scMeD&RNA-Seq

#### Barcode demultiplexing, read alignment, duplicate corrections, and DCM/CpG site identification

scMeD-Seq data was processed differently for plate based and combinatorial indexing approaches. To identify cells based on combinations of three barcodes, we generated all possible barcode combinations (i.e. virtual cells) *in silico*. If a barcode combination matched a barcode sequence at the beginning of a read in the FASTQ files, that read was annotated to the corresponding virtual cell. Once annotated, the barcode is trimmed from the read and downstream processing follows the same workflow as for plate-based scMeD-Seq.

For plate-based scMeD-Seq the first 20-23 nucleotides of the Read1 FASTQ files were trimmed using Cutadapt^59^ (-g AGTTCTACAGTCCGACGATC; v2.6). The trimmed reads were annotated based on their cell barcode, allowing 0 mismatches between the sequenced barcode and the reference barcode.

For both scMeD-Seq and sciMeD-Seq, the annotated reads were mapped to the mouse genome (mm10), using bowtie2^60^ (v2.4.1) local alignment (--local) and reads were separated based on CpG or DCM methylation. The reads were kept if an LpnPI cutting site (either CpG or DCM) was found inside or outside the read at a 12 to 16 nucleotide distance from the barcode sequence consistent with LpnPI generating a 32bp fragment from the center of the recognition site ^25^. Reads were annotated as methylated DCM sites if ‘CCAGG’ or ‘CCTGG’ sequences were found at the described position. Reads were annotated as methylated CpG sites if ‘CCG’, ‘CGG’, or ‘GCGC’ sequences matched the predicted location of LpnPI. The reads were processed to remove duplicates based on read sequence (combinatorial indexing) as well as UMI sequence (plate-based assays) and BAM files headers were modified making them suitable for SAMtools markdup –barcode-name (v1.18). Each unique DCM read contained a DCM site number, which was linked to the corresponding gene containing that site. These DCM site numbers were used to generate gene-based count matrices per barcode. This matrix was used for downstream analysis in R or python.

#### Resolving barcode combinations in sciMeD-Seq using combinatorial indexing

In all performed sciMeD-seq experiments, more unique barcode combinations than the number of expected nuclei were consistently observed. However, most of these barcodes yielded low read counts per barcode likely representing adapter ligation collisions, a pattern expected from combinatorial indexing techniques^57^ . To identify the barcodes that correspond to real nuclei, a cutoff based on a “Valley point” was applied in the barcode rank plot. This plot sorts barcodes by read count, with the valley representing the local minimum before the number of reads starts to rise again due to background noise or barcode collisions. This local minimum was calculated for each experiment, retaining barcodes above this cutoff. This number of barcode combinations retained typically fell in the same range as the number of nuclei in the experiment (∼500-1000 nuclei) that were manually counted prior library preparation (Figure S1B).

#### Barcode collision analysis

Expected barcode collision rate was estimated under a Poisson occupancy model. The probability that a given cell shares its barcode with at least one other cells is P(collision) = 1 – e^-n/b, where n are the distributed cells and b total available unique barcodes. For the combinatorial indexing experiments in small intestine, three rounds of barcode ligations yield b = 32^3 barcodes for the ∼750 cells that were sequenced resulting in an expected collision rate of 2.26%.

### Downstream data analyses of pre-processed sciMeD-Seq data

#### Count normalization and gene/cell filtering

Barcode count matrixes were generated and further processed by filtering out barcode combinations covering less than 200 genes. Depth/complexity outliers (n=1) (Tukey’s fence, k=1.5xIQR on log1p total counts or detected genes) were also removed. The depth-normalized, log-transformed counts for each cell were scaled to unit deviation and clipped (*scanpy.pp.scale*, max_value = 10), separately per batch and per assay to account for dynamic range differences.

#### BBKNN Integration with scRNA reference and dimensionality reduction

Following preprocessing, sciMeDseq data was combined with the reference scRNA dataset in scanpy, only including genes detected in both sets. For dimension reduction, batch-aware highly variable gene (HVG) selection was performed (*scanpy.pp.highly_variable_genes*, n_top_genes = 2,000, batch-aware across datasets), followed by principal component analysis (PCA) on the scaled, HVG-restricted expression matrix.

sciMeDseq and reference cells were integrated into a common UMAP embedding with BBKNN (*scanpy.external.pp.bbknn*, neighbors_within_batch = 1), swept across a range of dimensionalities (5–50 principal components) to assess the stability of the resulting embedding and cell-type recovery. At each dimensionality, Leiden community detection was performed at a fixed resolution (resolution = 1.0), and clusters were annotated by majority-vote label transfer from the reference dataset, and then collapsed to the bare cell-type label. Twenty principal components were selected as the working dimensionality for downstream analyses, producing results broadly consistent with the majority of other dimensions.

#### Gene set scoring (Module scores) of cell type marker gene sets

Cell type specific/enriched marker genes were obtained using scanpy.rank_genes_groups for each cell type versus all other cells in the reference dataset. In order to account for cell-to-cell heterogeneity, we utilized a t-test with conservative variance estimates (method = ‘t-test’). Genes were considered enriched in a cell type if log2(fold change) was greater than 1 and (Benjamini-Hochberg) adjusted p-value less than 0.0001, with genes being removed if they appear as significantly enriched in more than one cell type. Besides this differential expression analysis based on logistic regression was performed and genes with logreg score >0.5 and significant based on t-test DE analysis were kept together with known literature genes, making a gene set of 127 genes across 7 cell types.

The nonredundant lists of cell type enriched marker gene sets were then used to score reference and sciMeD-Seq cells for overall average expression per cell (*scanpy.tl.score_genes*, ctrl_size=50, n_bins = 25). Gene scoring was performed independently on each dataset, using the depth-normalized, log-transformed counts. Resulting gene set scores (module scores) per cell were further scaled to zero mean and unit deviation (per gene set, across cells) separately within each dataset, prior to Gaussian mixture modelling. cells for overall average expression per cell. Marker gene set enrichment was visualized using scaled gene set scores for each cell type enriched gene set across all cell types.

#### Gaussian mixture modelling for cell type classifications based on gene set enrichment

To classify sciMeDseq cells by cell type, we fit a joint Gaussian mixture model (GMM; scikit-learn GaussianMixture, n_components = 7) on the scaled marker gene set (module) scores of the query cells, across multiple random initializations. Fitted components were matched to reference cell types by solving a linear assignment problem that minimizes the Euclidean distance between component means and reference centroids in module-score space; each cell was assigned the cell type with the highest resulting posterior probability.

Classification confidence was controlled via a direct Bayesian false discovery rate (FDR) procedure (Newton et al., 2004): for each cell, a local FDR was defined as one minus its maximum posterior probability, and calls were retained if they fell within the largest ranked prefix whose running mean local FDR remained ≤5%, applied both globally and within each predicted cell type. Cells whose mixture log-density fell below the 1st percentile of the reference distribution were flagged as not well explained by any reference cell type. Call ambiguity was further assessed via the margin between a cell’s top two posterior probabilities (margin < 0.10 flagged as contested).

sciMeD-Seq cell annotation was based on two independent assignment methods: BBKNN integration and GMM. Cells with matching labels from both methods were used for further analysis.

#### Preprocessing of RNA data for scMeD&RNA-Seq

RNA reads were demultiplexed and converted to FASTQ files using bcl2fastq software (Illumina), as each cell contained unique Illumina indexes. For each single cell FASTQ file, the TSO primer sequence was removed, annotating the next 8 nucleotide UMI to the header of the read, using ‘umi_tools extract’ ^61^. The extracted reads were aligned to the reference genome downloaded from GENCODE (GRCm38.p4) using the default settings bowtie2 ^60^ (v2.4.1). SAMtools ^62^ (v1.18) was used to generate intermediate bam files for downstream processing analysis. The aligned reads were assigned to genes from GENCODE annotated mm10 GTF files, using *featureCounts* (Rsubread package ^63^, v2.0.1) at default parameters. Duplicate reads based on UMIs were removed and count matrixes were created using *umi_tools count* (parameters: --per-gene --gene-tag=XT --wide-format-cell-counts). The counts files per cell were merged, generating a count matrix for downstream analysis in R or python.

### Downstream analyses of pre-processed data

#### Analysis of DCM induction across genomic loci (genes)

Single cells were pseudobulked based on their dox condition and their DCM and CpG profile signals across genes were drawn to verify DCM-POLR2B activity. Genes were split in 100 bins, each bin corresponding to 1% of gene’s length, while upstream and downstream the TSS and TES corresponded to 100bp. DCM and CpG reads were normalized by their respective number of DCM and CpG sites per bin, respectively. To normalize for induction, the normalized signal was divided by the number of total DCM counts per condition. This was followed by visualization of DNA methylation signals depicting pseudobulk distributions.

#### Count normalization and gene/cell filtering

Raw count matrices for both DCM and mRNA assays were imported into scanpy (v1.11.5) for all cell barcodes identified. Cells with less than 200 detected genes (*scanpy.pp.filter_cells, min_genes = 200*), and genes which were detected in less than 3 cells (*scanpy.pp.filter_genes, min_cells = 3*) were removed from further analysis. Raw counts were depth-normalized per 10,000 unique transcripts (*scanpy.pp.normalize_total*, target_sum = 1e4) and log-transformed (*scanpy.pp.log1p*) for each cell. The depth-normalized, log-transformed counts for each cell were further scaled to unit deviation and clipped (*scanpy.pp.scale*, max_value = 10), separately per batch and per assay to account for dynamic range differences.

#### Dimensionality reduction and integration with reference scRNAseq datasets

Previously published data for small intestine scRNAseq reference were obtained as raw count matrices and cell metadata from GSE92332. Raw scRNAseq count matrices were processed as for scMeD&RNA above (section: ‘*Count normalization and gene/cell filtering*’). Following preprocessing, the reference dataset was combined with our scMeD&RNA dataset in scanpy, keeping only genes detected in all datasets (*anndata.concatenate*, join = ‘inner’). For dimension reduction, batch-aware highly variable gene (HVG) selection was performed *(scanpy.pp.highly_variable_genes*, min_disp = 0.5, min_mean = 0.0125, max_mean = 3, n_top_genes = 2,000) followed by principal component analysis (PCA) on the HVGs.

All scMeD&RNA and reference cells were integrated into a common UMAP embedding with BBKNN (v1.6.0) using the top 25 PCA components identified (*scanpy.external.bbknn*, n_pcs = 25, neighbors_within_batch = 1). Clustering was then performed using leiden community detection under default settings (resolution = 1), and clusters were automatically annotated by cell type label transfer from the reference dataset based on each cluster’s majority label. Leiden clusters were collapsed to major cell type, in order to obtain similar annotation resolution to the reference dataset. To verify robustness of integration, Pearson correlation matrices were quantified between sMeD&RNA and reference cells (*scanpy.tl.dendrogram*, method = ‘pearson’) in PCA latent space.

#### Gene set scoring (Module scores) of cell type marker gene sets

Cell type specific/enriched marker genes were obtained by scanpy.rank_genes_groups for each cell type versus the rest in the reference dataset. In order to account for cell-to-cell heterogeneity, we utilized a t-test with conservative variance estimates (method = ‘t-test’). Genes were considered enriched in a cell type if log2(fold change) was greater than 1 and (Benjamini-Hochberg) adjusted p-value less than 0.0001, with genes being removed if they appear as significantly enriched in more than one cell type.

The nonredundant lists of cell type enriched marker gene sets were then used to score scMed&RNA cells for overall average expression per cell. Gene scoring was performed on the depth-normalized, and log-transformed counts versus a random background of 50 genes per expression bin (*scanpy.score_genes*, ctrl_as_ref = True, ctrl_size = 50, n_bins = 25). Resulting gene set scores (Module scores) per cell were further scaled to unit deviation (per gene set, between cells) separately per batch/assay prior to Gaussian mixture modelling. Marker gene set enrichment was visualized using scaled gene set scores for each cell type enriched gene set across all cell types.

#### Gaussian mixture modelling for cell type classifications based on gene set enrichment of DCM

In order to distinguish active (matching) versus inactive (mismatching) cells for a given cell type enriched gene set, we utilized a Gaussian mixture model (GMM) on the scaled marker gene set scores calculated above. GMM was performed using the GaussianMixture model from scikit-learn (v1.8.0) with n_components = 2 (active/inactive). The resulting posterior probabilities - which indicate the probability that a given cell belongs to the active (matching) population - were used to classify cells given that a posterior threshold greater than 0.85 is met for a given cell. Cells were annotated and kept based on GMM calling if the posterior threshold for only one gene set is met. GMM modelling was also performed separately per sequencing batch and per assay to account for batch effects/dynamic range differences between assays.

#### Annotating single-cell state transitions based on past and present gene expression

To identify the bona fide transitions, we used DCM cell labels based on the GMM analysis (FDR<0.05) reflecting past cell state while the RNA profile (present state) was based on BBKNN integration as it showed strong positive correlation with reference data (Fig. S5H). Multiple labels per cell were allowed for DCM data providing p-value thresholds were met. To measure cell state stability across modalities (pulse / chase), cells were grouped accordingly and divided between a number of stable (labels matching) and unstable labels mismatching) states the latter indicating transitions.

### Predicting future cell fate decisions using CellRank2

CellRank2 was used to predict lineage dynamics across clusters within the integrated cell embeddings^43^. To generate input data for RNA velocity analysis, BAM files from the reference dataset were downloaded^31^, and unique pRNA reads were processed. These reads were split into intronic and exonic counts using *velocyto run_smartseq2* (or *velocyto run*), producing *.loom* files compatible with downstream velocity analysis^39^. During preprocessing, reads overlapping repetitive regions (as defined by the mm10_rmsk.gtf annotation from UCSC) were excluded to ensure accurate quantification.

The loom files for each batch and dataset were merged and the cells containing RNA used in previous integration and downstream analyses were included, excluding other cells. UMAP and PCA coordinates from the previous integration (see above) were transferred and CellRank2 analysis was performed. Scanpy was used for processing the count matrix. Genes with less than 3 counts across all cells were discarded and counts were log normalized and scaled to a max value of 10. For RNA velocity genes with minimum 20 counts across all cells were included and the top 3000 highly variable genes were filtered. The neighborhood graph was computed using scanpy.pp.neighbors (n_neighbors = 30 and n_pcs =30) based on the top principal components. To prepare for RNA velocity estimation, scvelo.pp.moments was applied (n_pcs=None, and n_neighbors=None) reusing the computed neighborhood graph. RNA dynamics were estimated using the dynamical model implemented in scVelo^40^. To model cell state transitions, *VelocityKernel* was constructed and its transition matrix computed using the *compute_transition_matrix* function. This matrix was then passed to the GPCCA estimator (*cellrank.estimarors.GPCCA*) to decompose the transition matrix into macrostates. To ensure representation of all defined clusters, 25 macrostates were computed arbitrarily using the *compute_macrostates* function, with the goal of capturing at least one macrostate per cluster. All macrostates were treated as terminal states, while only ISCs were defined as the initial state.

Fate probabilities were calculated using *compute_fate_probabilities* function. The computed fate probabilities for each macrostate were aggregated by the cluster of origin, resulting in a cluster-to-macrostate fate probability matrix that summarizes the average differentiation potential of each cluster toward the inferred terminal states. Individual cells that showed reversion toward the ISC state were assigned to distinct clusters, enabling the quantification of their fate probabilities toward each inferred macrostate.

For CytoTRACE modality all counts were used, and CytoTRACEKernel was used to model cell state transitions^41^. The transition matrix was computed with default settings (threshold_scheme=”soft”, and nu=0.5). The remaining analysis steps were performed as described for *VelocityKernel*.

For pseudotime analysis we used the diffusion pseudotime (dpt) package^42^. Expression values were log normalized followed by computation of diffusion maps (*scanpy.tl.diffmap*). The starting state was designated as the root index based on its position within the diffusion map corresponding to the ISC and TA clusters. Terminal states were identified as cells located at the greatest diffusion distance from this initial state, representing the endpoints of the differentiation trajectory. Dpt pseudotime was computed by running *sc.tl.dpt* with default settings. Dpt pseudotime was incorporated into the PseudotimeKernel to compute the transition matrix and subsequent fate probabilities, following the same approach described above.

A combined transition kernel was constructed as a weighted sum of individual kernels to capture multiple aspects of cell state transitions. Specifically, the combined kernel was calculated as: *velocity *0.3 + dpt*0.3 + cytotrace*0.3 + connectivity*0.1* where *velocity* represents the RNA velocity kernel, *dpt* the dpt pseudotime kernel, *cytotrace* the connectivity kernel, and connectivity the velocity connectivity kernel. The weights reflect the relative contribution of each kernel in modeling cell fate transitions.

### Single cell state tracing in Neural differentiation

#### Mouse embryonic stem cell (mESC) culture

To obtain feeder cells, DR4 Female Mouse Embryonic Fibroblasts (MEFS, Passage 1, C57BL/6)^64^ were cultured in MEF medium: Dulbecco’s Modified Eagle Medium (DMEM) (Gibco; 41966-029), supplemented with 15% Fetal Bovine Serum (FBS) (Capricorn Scientific; FBS-12A), 100 U/mL Penicillin and 0.1 mg/mL Streptomycin (Sigma; P0781), 1X Non-Essential Amino Acids (NEAA) (Gibco; 11140-050), 0.1 mM 2-mercapto-ethanol (Gibco; 31350-010) and plated on 15cm non-coated plates at 50% confluency. Cells were split every 2-4 days at 80-100% confluency to expand these by using trypsin-EDTA (Sigma; T3924). MEFs were collected and irradiated at 195 kV and 10.0 mA for 30 minutes using a GammaCell Irradiator and frozen in MEF medium and DMEM supplemented with 20% DMSO and 20% FBS at a 1:1 ratio.

Mouse embryonic stem cells (mESCs) containing the *dcm-Polr2b-Fkbp12-cre* fusion- gene (mESC^dcm-pol-fkbp12-cre^) were cultured at 37°C with 5% CO_2_ on 0.2% gelatin-coated plates seeded with a feeder layer of gamma-irradiated MEFs in ES medium (Dulbecco’s Modified Eagle Medium (DMEM) (Gibco; 41966-029), 15% Fetal Bovine Serum (FBS) (Capricorn Scientific; FBS-12A), 100 U/mL Penicillin and 0.1 mg/mL Streptomycin (Sigma-Aldrich; P0781), 1X Non-Essential Amino Acids (NEAA) (Gibco; 11140-050), 0.1 mM 2-mercapto- ethanol (Gibco; 31350-010) and 1X Leukemia Inhibitor Factor (LIF)). 3 µM CHIR99021 (Sigma; SML1046) and 1 µM PD0325901 (Selleckchem; S1036) were added to ES medium prior to use. Cells were passaged every 2-3 days using trypsin-EDTA (Sigma; T3924), maintaining a density between 0.2×10^6^ and 1×10^6^ cells per mL in wells of a 6-well plate.

#### Design of the DcmPol2B-fkbp12-cre system

As the conventual DCM-Pol2B system in the collagen locus does not express in all cell types including the neural and hematopoietic lineages, a more versatile DcmPol2B-fkbp12-cre construct was synthesized using GenScript Gene Synthesis services (fig. S8A) for targeting into the Tigre locus. Compared to the original system, the fusion protein contains a Flag-tag for easier detection and a fkbp12 degradation tag enabling rapid degradation of the fusion protein using dTAG-13^65^. A Cre protein was added and expressed using a P2A sequence following the fusion protein, which will remove a downstream stop signal flanked by loxP sites. This enables an opposing PGK promoter to express *mecfp*, in the opposite direction further interfering with fusion protein expression and irreversibly marking induced cells for identification. The construct contains PiggyBac elements flanking the whole construct allowing for random integration, while the insulators (IS) avoid silencing or inappropriate activation. An FRT element is located downstream of a PGK promoter allowing for hygromycin selection upon integration.

Tigre homology arms, Ori site and Ampr sequences were amplified from the Tigre plasmid (gift from [Neil Brockdorff](https://www.nature.com/articles/s41467-019-11171-3#auth-Neil-Brockdorff)) ^66^ using primers containing restriction enzyme (RE) overhangs (Table S4, PCR cloning primers) to enable targeting to this locus. The Col1a1-frt-hygro-pA (Addgene, 20730) was digested with NotI and BstEII RE, and the frt-Neo^R^-frt-hyg^R^ fragment was isolated. The amplified Tigre fragment and isolated selection cassette were ligated and transformed into NEB® 5-alpha Competent E. coli (NEB, C2987H), generating the intermediate Tigre-frt-hygro-pA plasmid. Subsequently, the PGK promoter and rTTA sequences were amplified (Table S4, PCR cloning primers) and cloned into the linearized Tigre-frt-hygro-pA, which had been digested with BstEII-HF. Assembly of the PGK and rTTA inserts with the linearized vector was performed using Gibson assembly following the manufacturer’s protocol, to generate the final Tigre-frt-hygro-pA-PGK-rTTA plasmid. The plasmids were expanded using QIAGEN Plasmid Maxi kit (Qiagen, 12162) according to manufacturer’s instructions.

#### Targeting the DcmPol2B-fkbp12-cre system to the TIGRE locus in mESCs

Castaneus mESCs were cultured as described above. Cells were transfected with lipofectamine 2000 (ThermoFisher, 11668019), following manufacturer instructions. The Tigre-frt-hygro-pA-PGK-rTTA plasmid was integrated into the Tigre locus using pSpCas9(BB)-2A-Puro (PX459) V2.0 (Table S4. gRNA) (Addgene, 62988). After 2 days of 140µg/mL puromycin (Sigma, P4512) selection, colonies were picked and transferred to a 96 well plate for expansion and genotyping using primers targeting for DCM-Polr2b (Table S4. PCR genotyping). Colonies with correct integration were further expanded and characterized for rTTA expression. The newly generated mESC^frt-hyg-rtta^ were used for integration of the DcmPol2B-fkbp12-cre system. Lipofectamine 2000 was used as described previously, together with the DMCPOL-fkbp12-CRE plasmid, pCAGGS-flpE-puro (Addgene, #20733) and GFP reporter (pLKO5.sgRNA.EFS.GFP, Addgene, 57822). After 24h, GFP-positive cells were FACS sorted into 10 cm plates containing ES medium with Puromycin and Hygromycin (InvivoGen, ant-hg-1) selection. After 1-week colonies were picked for expansion and genotyped for correct integration of the DCM construct (Table S4, PCR genotyping). Clones containing correct integrations were further expanded and characterized for DMCPOL-fkbp12- CRE and mECFP expression upon dox induction (fig. S7B). Expression of the fusion protein was induced using 2µg/µL doxycycline dissolved in water, and protein destruction was initiated by adding 500nM dTAG-13 (Tocris, 6605) dissolved in DMSO (Sigma, D2438). This line is further referred to as mESC^dcm-pol-fkbp12-cre^.

#### Monitoring Pol2B-DCM induction in mESC^dcm-pol-fkbp12-cre^ using qPCR

Following dox induction (2µg/mL) cells were collected for subsequent RNA isolation using the ReliaPrep™ RNA Cell Miniprep System according to the manufacturer’s instructions (Promega, #Z6010). Reverse transcription was performed using 1µg of the isolated RNA, 1µL oligo-dT (Thermo Fisher, SO132) and SuperScript Reverse Transcriptase III (Thermo Fisher, 18080044). Quantitative PCR was performed using 20x diluted cDNA and following GoTaq qPCR MasterMix Protocol (Promega, A6001). The qPCR was performed according to the following protocol in CFX Opus Real-Time PCR Systems (Bio-Rad): 1 cycle of 95°C for 2 minutes, 40 cycles of 95°C for 30 seconds, 60°C for 30 seconds, 72°C for 30 seconds, 1 cycle of 72°C for 1 minute. *Gapdh* was used as reference control and relative expression was calculated by the 2-^ΔΔCt^ method. The primers used in this experiment are described in Table S4.

#### Monitoring Pol2B-DCM induction in mESC^dcm-pol-fkbp12-cre^ using western blot analysis

To create nuclear extracts (NPX) for western blot analysis, cultured cells were collected in 1mL PBS + 1x Protease inhibitor (PI; Roche, 11873580001) in in 1.5 ml Eppendorf tubes. The cells were centrifuged at 300g for 5 minutes and the supernatant was removed. The pellet was resuspended in 400µL buffer A (10 mM Hepes pH 7.6, 1.5 mM MgCl_2_, and 10 mM KCl) by flicking the tube, followed by 10 minutes incubation on ice. Right after, the sample was vortexed for 30 seconds, and centrifuged at 1,600g for 5 minutes at 4°C.The supernatant containing cytoplasmic proteins was stored for analysis. The pellet, containing nuclear proteins, was resuspended in buffer C (20 mM Hepes pH 7.6 (Gibco, 15630056), 25% glycerol (Sigma, G9012), 420mM NaCl, 1.5 mM MgCl_2_, and 0.2 mM EDTA) by flicking the tube. The sample was incubated on ice for 20 minutes with regular tube flicking. Samples were centrifuged at max speed (17,000g) for 10 minutes at 4°C and the supernatant containing the NPX was transferred to a new tube. The concentration of all samples was measured using a DeNovix spectrophotometer (DeNovix) and concentrations were equalized between samples using buffer C.

The NPXs were combined with LDS buffer mix (4x LDS (Invitrogen, NP0007) and 10x reducing agent (Invitrogen, NP0009) to a final 1x concentration). A total of 80µg of protein per sample was boiled at 95°C for 5 minutes, vortexed and kept at room temperature. The Criterion TGX Precast 4-15% polyacrylamide gel (Bio-Rad, 5671084) was used with 1X Tris-Acetate SDS running buffer pH8.3 (Bio-Rad, 161-0772), together with 5µL protein ladder Precision Plus Protein Dual Color Standards (Bio-Rad, #61-0374). The gel was run at 100V for 15 minutes followed by an increase to 150V for 45 minutes. The transfer was performed with the Bio-Rad Turbo blotter following the high MW protocol of 10 minutes into 0.2µm PVDF membrane (Bio-Rad, 1704157). Membranes were blocked for 1 h at room temperature in 2.6% skim milk powder (Sigma-Aldrich, 70166) dissolved in 1x TBS buffer (25 mM Tris-HCl, 150 mM NaCl, pH 7.6). After blocking, the membrane was incubated with primary antibody in blocking buffer. To target DCMPOL2, mouse anti-V5 (Invitrogen, R96025, 1:5,000) was used for 2h at RT. After three washes with TBST washing buffer (1x TBS, 0.1% Tween-20), the membrane was incubated with secondary goat anti-mouse IgG-peroxidase antibody (Sigma, A4416; 1:5,000) for 1 h at room temperature in blocking buffer supplemented with 0.1% Tween-20 (Sigma, P9416). As an internal protein control, β-actin was used: mouse monoclonal anti-β-actin-peroxidase antibody (Sigma, A3854, 1:10,000). Protein content was determined with SuperSignal West Femto Maximum Sensitivity Substrate (Thermo Fisher, 34094) analyzed using an Amersham Imager 600 (Amersham Biosciences).

#### Analysis of DCM enrichment at non-coding DNA in mESCs

Deeptools (v3.5.1) ^67^ was used to generate metagene plots of pseudobulk DCM reads, on different genomic regions: gene bodies and mESC histone modifications (H3K27Ac, H3K4me3, H3K36me3, H3K27me3 and H3K9me3) ^68^. Single cell data was merged in pseudobulk samples and downstream analysis was performed as shown previously for bulk data ^25^. Briefly, intragenic distribution of DCM (or CpG) was plotted by generating 100 bins from 10kb either upstream of the TSS or downstream of the TES. Gene body bins were generated by adding 1% of the gene size into each bin, generating a total of 100 bins. The signal per bin was adjusted for DCM or CpG sites. Different gene sets reflecting low or high expressing genes were classified based on external mESC RNA expression data ^25^.

Enriched region files for each histone modification were downloaded and used to intersect datasets. Reference data for histone modifications in mESCs were downloaded from different sources: mESCs histone modifications (ES-Bruce4) were downloaded from Encode^68^: H3K4me3 (ENCSR000CBG), H3K4me1 (ENCSR000CBF), H3K27ac (ENCSR000CDE), H3K36me3 (ENCSR000CFO), and H3K9me3 (ENCSR000CFZ); EnhancerAtlas2.0 (mm9)^69^: ESC_R1, ESC_KH2, ESC_J1, ESC_D3, ESC_D0, ESC_C6, ESC_Bruce4, and ESC_46C; mm10 annotation (Gencode): GRCm38.p4; mESC bulk RNAseq from: SRX8023826, SRX8023827 and SRX8023828. mm9 regions were converted to mm10 regions using the liftOver function from UCSC^70^. Unique DCM and CpG reads were normalized by using BamCoverage (--binSize 25 --centerReads --ignoreDuplicates --scale factor 0.598 or 1 for +/- dox conditions, respectively). The scale factor refers to the ration of DCM versus CpG read differences between induced and non-induced conditions. BigWig output files were combined with BED files for the different histone modifications to compute the scores per region using computeMatrix (scale-regions -m 1000 -b 4000 -a 4000). plotHeatmap or plotProfiles were used to visualize the data. Genome tracks were visualized using Integrative Genomics Viewer (IGV), by plotting normalized BigWiG files.

#### Motif Enrichment analysis

To predict TFs binding sites from induced DCM site regions, the captured DCM sites were converted to fragment files with counts per region and 250bp were extended upstream and downstream the DCM sites. mESC enhancer bed files were obtained from the EnhancerAtlas2.0 ^69^, merged, and overlapped with extended DCM regions. The intersected regions were analyzed using Homer software (findGenomeRegions.pl function with the following settings (-size 200, -bg (-dox sample, -MaskMotif LpnPI.motifs)) by using LpnPI motifs as motif background and -dox regions to reduce false positives. The top 100 motifs log10 P-value were plotted.

#### Differentiation of Embryonic Stem Cells (ESCs) into Neural Progenitor Cells (NPCs)

mESCs^dcm-pol-fkbp12-cre^ were collected and re-plated on non- coated 10cm plates for 15 minutes in ES medium to remove MEF feeder cells. After incubation, floating mESCs^dcm-pol-fkbp12-cre^ cells were collected, washed twice with5 mL N2B27 medium (spun at 300g) and plated on gelatin-coated plates to induce neural differentiation. N2B27 medium was prepared using [1:1] DMEM/F12 (Gibco; 31330-038) and Neurobasal (Gibco; 21103-049), 2 mM L-glutamine (Gibco; 25030-024), 1X N2 (Gibco; 17502-048), 0.5X B27 (Gibco; 17504-044), 0.1 mM 2- mercapto-ethanol (Gibco; 31350-010) and 100 U/mL Penicillin and 0.1 mg/mL Streptomycin (Sigma-Aldrich; P0781). Medium was refreshed every other day for 7 days. On day 7, cells were detached by using StemPro™ Accutase™ Cell Dissociation Reagent (Gibco; A1110501) and plated at 3×10^5^ cells in non-gelatinized 6-well plates in 2mL N2B27 medium supplemented with 10 ng/mL EGF (Peprotech; 315-09) and 10 ng/mL FGF (Peprotech; 100-18B) while shacking at 75 rpm until day 10. On day 10, embryoid bodies (EBs), formed during day 7-10, were collected by centrifugation of medium at 150g for 3 minutes and EBs were plated into 1% matrigel (Corning; 354277) coated wells in N2B27 medium + 10ng/mL EGF and FGF. On day 14, NPCs were plated at a density of 2×10^5^ cells per mL to remove remaining undifferentiated cells and EBs.

NPCs were further differentiated following a protocol to generate mixed neural populations^46^. Briefly, the NPCs were seeded in Matrigel coated plates at 135,000 cells per well of a 6-well plate. The next day, the medium was changed to D1 medium (Euromed-N (Euroclone, ECM0883L), 1% B27, 0.5% N2, 1% penicillin streptomycin (P/S)), supplemented with 10ng/mL FGF. After 3 days, medium was changed to medium A (1:3 DMEM/F12 and neurobasal medium, 1% B27, 0.5% N2, 1% P/S), supplemented with 10ng/mL FGF and 20ng/mL BDNF (Peprotech, AF-450-02). After an additional 3 days the medium was changed to medium B1 (1:3 DMEM/F12 and neurobasal medium, 1% B27, 0.5% N2, 1% P/S) supplemented with 6.7 ng/mL FGF and 30 ng/mL BDNF.

#### Bulk MeD-Seq sample preparation of induced NPCs

Cells were collected and gDNA isolation was performed using the QIAamp DNA micro kit (Qiagen, #56304). MeD-seq sample preparation was performed as previously described^28^. Briefly, gDNA samples were isolated using QIAamp DNA micro kit (Qiagen, #56304) and digested by LpnPI (New England Biolabs, R0663S). Stem-loop adapters were ligated to repaired DNA and amplified with Illumina dual indexed barcodes using high-fidelity polymerase. Amplified products were size-selected using a Pippin HT system (Sage Science) with 3% agarose cassettes (Sage Science, HTG3010), then multiplexed samples were sequenced on Illumina HiSeq2500 (50bp single reads) and demultiplexed using bcl2fastq.

#### Bulk MeDseq data analysis of induced NPCs

Raw data was processed as shown previously ^25^. Illumina adaptors were trimmed from the raw FASTQ reads, followed by a filtering step for fragments containing an LpnPI restriction site between 13bp and 17bp from the 5’ or the 3’ end of each read. Reads directly linked to DCM methylation and CpG methylation were separated and mapped to the mouse genome (mm10) using bowtie2 ^60^. Annotated genes from UCSC (GRCm38.p2) were used to generate DCM read scores. SAMtools version 0.1.18 was used to create BAM files for visualization in IGV ^62,71^. For absolute DCM enrichment, DCM read coverage was normalized to CpG read coverage from the same sample. For relative DCM enrichment, DCM coverage was normalized between induced samples. In both cases, read counts were normalized per million reads sequenced (RPM). Absolute DCM levels refer to the efficiency of DCM-Pol2rb induction, while relative DCM levels can be used to normalize for induction differences between samples/time-points.

DCM-POL2RB expression was induced in NPCs^Tigre(dcm-pol-fkbp12-cre)^, by adding dox for 16h at day 2 and refreshing medium with 500nM dTAG-13 following induction to immediately degrade DCM-POL2RB protein. dTAG-13 medium was kept on the cells for a chase of 48h. Control samples such as +dox 16h of day 2 and day 4, -dox day 2 and day 4 and DMSO chase (no-degradation) were also generated. To identify methylation differences between using dTAG or DMSO, several analysis following gene filtering were performed: 1) Genes below the DCM induction threshold were filtered out. Differential expression analysis was performed between DCM counts of non-induced and induced samples. In this analysis, the size factors were replaced for the total CpG reads of each sample. Genes above and below log2(FC) 1 and -1, and FDR < 0.05 were identified as induced genes. 2) Differentially methylated genes (DMGs) were calculated between induced samples: +dox day 2 and day 4, and +dox day 2 followed by a chase to day 4 with dTAG or DMSO. The DMGs log2(FC) of +dox day 2 vs +dox day 4 were plotted against +dox day 2 vs chase dTAG or chase DMSO (fig. S7F). To compare induced samples, the size factors were replaced for the total DCM reads of each sample. These analyses were performed using Deseq2 (v1.44).

#### Single Cell RNAseq to create a reference dataset for neural differentiation

Cells from different samples were collected simultaneously and processed using 3’CellPlex kit set A (10x Genomics, PN-1000261). Briefly, the cells were collected and resuspended in 1mL buffer (PBS + 10% FBS). After centrifuging for 5 minutes at 300g, the cell pellets were resuspended in 100µL of CellPlex oligo. The oligo contains a lipid that binds to the membrane, tagging the cells. After 5 minutes incubation, the samples were washed with 1.9mL buffer (1% BSA in PBS) and centrifuged for 5 minutes at 300g. Cells were resuspended and counted with the Countess II (Life Technologies). Different samples were pooled at equal numbers of live cells. Mixed cells were stained with a 1:10,000 dilution of 10mg/mL Hoechst (Molecular Probes, H3570), and live cells were sorted using FACS AriaII cell sorter (BD Biosciences) and centrifuged for 5 minutes at 300g. Pooled cells were resuspended in PBS + 10% FBS at a concentration of 1,400 cells/µL. The samples were processed using Chromium Next GEM Single Cell 3’ v3.1 (10x Genomics, 1000269). The CMO library was prepared using 3’ Feature Barcode Kit (10x Genomics, P-1000262) to amplify the conjugated oligos and the dual index kit NN (10x Genomics, 1000243) for library indexing. 10x Genomics dual indexing parameters were: 28/10/10/90 cycles for R1/i7/i5/R2 reads. Libraries were sequenced on the Novaseq 6000 platform.

#### Single cell Multiome (ATAC-Seq + RNA-Seq) sample processing to generate a reference dataset for neural differentiation

##### Dead cell removal and nuclear isolation

Dead cells were removed using the dead cell removal kit (Miltenyi Biotec, 130-090101). Briefly, the collected cells were resuspended in 10µL beads per 1 million cells and incubated for 15 minutes at room temperature. Volumes were equalized at 500µL in 1x binding buffer, and the dead-lived cell separation was performed following the manufacturer instructions using LS columns (Miltenyi Biotec, 130-042-401). After dead cell removal, live cells were collected and lysed for 5 minutes on ice with lysis buffer (10mM Tris-HCl pH 7.4, 10mM NaCl, 3mM MgCl_2_, 1%BSA, 0.1% Tween-20, 1mM DTT and 1U/µl RNAse inhibitor) to isolate nuclei. After incubation, all samples were washed 3X with 1mL wash buffer (10mM Tris-HCl pH7.4; 10mM NaCl; 3mM MgCl2; 1%BSA; 0.1% Tween-20; 1mM DTT; RNAse inhibitor), spun down at 500g for 5 min, and nuclei were resuspended in 10µL nuclei buffer (10x Genomics, PN-2000153), counted using the Countess II (Life Technologies) and diluted to a final concentration of 10,000 nuclei/µL. For NPC differentiation, day 2 and day 4 samples were mixed at a 1:4 ratio as day 2 cells were more homogeneous.

##### Library preparation

Sixteen thousand nuclei from the mixed sample were used to conduct the Chromium Next GEM Single-Cell Multiome ATAC-seq and Gene Expression protocol (10x Genomics, PN- 1000280) according to the manufacturer’s instructions. Resulting libraries were quantified using qPCR, and fragment sizes were determined by bioanalyzer using a high sensitivity chip (Agilent, 5067-4626). The ATAC library was sequenced following paired-end dual indexing parameters: 50/8/24/49 cycles for R1/i7/i5/R2. All libraries were sequenced using the Novaseq 6000 platform (Illumina).

#### Preprocessing of scRNAseq (10x Multiplex) and multiome (GEX+ATAC) reference data for NPC differentiation

To generate a reference dataset for state tracing experiments in neural progenitors 10x and multiome data was generated. scRNAseq binary base call (bcl) data was transformed to FASTQ files using the *cellranger mkfastq* pipeline. To process scRNAseq multiplex samples, *CellRanger multi* was used with cell multiplex oligo (CMO) sequence references to assign the cells to their corresponding sample. Multiome samples were processed by using the Cellranger-arc tool. GEX and ATAC reads were aligned and quantified simultaneously using *cellranger-arc count*. Mouse samples for NPC differentiation were separated from other human samples that were mixed in, by aligning to a the mouse (mm10) and human (hg38) genome reference, downloaded from GENCODE databases (GRCm38.p4 and (GRCh38.p14)). Cells were assigned to human, or mouse based on their alignment scores (>80%). Reads were realigned to the mouse (mm10) reference genome and expression matrixes were obtained using the default parameters of CellRanger.

#### Data analysis of 10x scRNAseq data for NPC differentiation

##### Normalization, Clustering and UMAP

Count matrixes were loaded and cells with less than 500 UMI counts, 200 genes and more than 10% mitochondrial UMI percentage and 20,000 UMI counts were filtered out in R software^72^. The four time points of NPC differentiation (D0, D2, D4 and D6) were merged without integration as the CellPlex approach minimizes batch effects. Data was normalized using logarithmic normalization. The 2000 highest variable genes were selected using the vst method (within Seurat pipeline) and used to perform Principal component analysis using Seurat package. Elbowplot was used to determine the number of dimensions per dataset. Dimensionality reduction was performed using UMAP (dim=15), followed by the *FindClusters* function (resolution = 0.2). Marker genes per cluster were discovered using ‘FindAllMarkers’’ next parameters: min.pct = 0.25, logfc.threshold = 1, test.use = “wilcox”, Bonferroni correction is set as default to generate adjusted p-values.

##### Cluster annotation

The generated unsupervised clusters were manually annotated using known neural differentiation markers^46,73^. *Nestin, Hes5,* and *Slc1a3* genes were used to identify NPCs/RGCs; *Gria2, Tubb3, Dcx* and *Sox6* for immature dorsal neurons; and *Col3a1, Fn1, Prrx1* and *Twist1* for immature glial cells.

##### DCM in ATAC peaks

scATAC-seq peaks were first generated using the Cell Ranger ATAC pipeline, which provides high-confidence accessible chromatin regions across single cells. To quantify DCM sites within these regulatory regions, the ATAC peak regions were overlapped with DCM reads on a per-cell basis. This overlap analysis yielded a matrix of DCM read counts per peak per cell, enabling the construction of a DCM-in-ATAC-peaks count matrix.

##### Dimensionality reduction and Harmony Integration analysis

Reference datasets (10x chromium and multiome) together with pRNA and pDCM datasets (from day 2 and day 4) were processed, retaining cells with a minimum of 200 detected genes. Following preprocessing, all datasets were combined in scanpy keeping only genes detected across all datasets (sc.concat, join = ’inner’). For dimensionality reduction, batch-aware highly variable gene selection was performed across dataset batches (scanpy.pp.highly_variable_genes, n_top_genes = 3,000, batch_key = ’assay’), followed by scaling (max_value = 10) and principal component analysis on the selected HVGs (n_comps = 50). All datasets were integrated into a common embedding using Harmony (harmonypy, random_state = 42, max_iter_harmony = 20) on the top 20 PCA components, correcting for dataset of origin (‘assay’). A shared neighborhood graph was then constructed on the Harmony-corrected embedding (scanpy.pp.neighbors, n_neighbors = 30, n_pcs = 20), followed by UMAP visualization and leiden clustering (resolution = 0.1, random_state = 0). Clusters were labelled based on the previously defined gene markers.

##### State transitions across clusters between pDCM and pRNA

Putative state transitions were based on mismatches between unsupervised cluster assignments for DCM and RNA data from the same cell. To refine this analysis a module score was calculated followed by GMM analysis in a similar manner as described for the intestine. For the GMM analysis, a threshold of 0.837 was set to obtain a Bayesian false discovery rate < 0.05. The generated Z scores were visualized per cluster and a pairwise t- test was applied to verify score specificity. The DCM and RNA scores between RGCs, Ins and IGs were scaled across gene modules for visualization on a heatmap to compare transitions.

#### Statistical analysis

We used ggpubr (v0.6.0), stats (v4.4.2), scipy (v1.16.3) and packages with their own integrated stats such as the scanpy (v1.11.5) Python package to implement pairwise and two-sided independent samples t-test, binomial tests, Pearson correlations, statistics and associated P values used in this study. Scikit-learn (v1.8.0) was used for Gaussian Mixture Modeling and Plots were generated using ggplot2 (v3.5.1), or matplotlib (v3.10.3).

## Acknowledgments

M.P.C, G.V and F.F. were supported by KWF grant nr. 2021-2/14200. We thank Riccardo Fodde for critical reading of the manuscript. We thank Life Science Editors for editorial support.

## End notes

### Author contributions

E.G.S., F.F., G.V., M.P.C., designed experiments E.G.S., F.F., G.V., E.B, M.L. and A.S. performed experiments. E.G.S., J.K., R.B., J.B., and B.T. participated in formal data analysis. E.G.S., and M.P.C. interpretated the data. M.P.C. and J.G. conceived the study. M.P.C. and J.G. supervised the study and secured funding. E.G.S., and M.P.C. wrote the manuscript. All authors reviewed and edited the manuscript.

### Data and materials availability

All raw FASTQ data files, processed gene-by-cells count matrixes and metadata generated for this study are available upon publication at the NIH Gene Expression Omnibus (GEO) under accession number: GSE303518

Reference data for histone modifications in mESCs were downloaded from different sources: mESCs histone modifications (ES-Bruce4) were downloaded from Encode ^68^: H3K4me3 (ENCSR000CBG), H3K4me1 (ENCSR000CBF), H3K27ac (ENCSR000CDE), H3K36me3 (ENCSR000CFO), and H3K9me3 (ENCSR000CFZ); Enhancers for motif analysis were extracted from EnhancerAtlas2.0 ^69^ (mm9): ESC_R1, ESC_KH2, ESC_J1, ESC_D3, ESC_D0, ESC_C6, ESC_Bruce4, and ESC_46C; mm10 annotation (Gencode): GRCm38.p4; mESC bulk RNAseq from Boers et al.,: SRX8023826, SRX8023827 and SRX8023828 ^25^ . scRNAseq intestine data ^31^: Atlas and LargeCellSort UMI counts were downloaded from GSE92332. The bam files for each batch were downloaded to generate exonic and intronic counts (SAMN07976239-SAMN07976241, SAMN07976259-SAMN07976265; SAMN07958121- SAMN07958126 and SAMN07958175).

Supplementary Information is available for this paper.

**Materials & Correspondence** requests for materials should be addressed to

