## Supplementary figures for "Time-resolved single-cell state tracing exposes bidirectional state dynamics during differentiation"

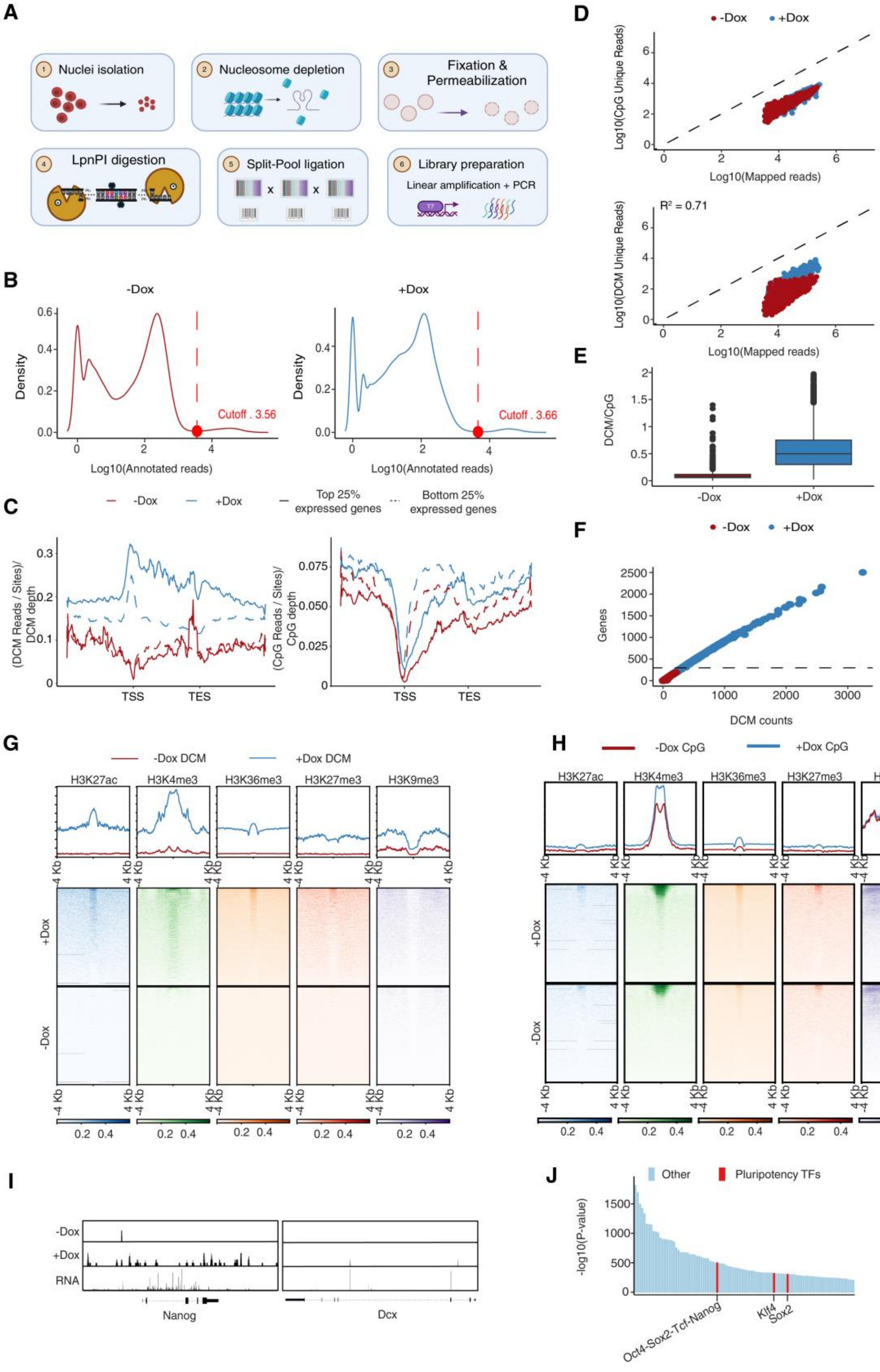
A
Nuclei isolation
Nucleosome depletion
Fixation & Permeabilization
LpnPI digestion
Split-Pool ligation
Library preparation
Linear amplification + PCR
B
-Dox
+Dox
Density
Log10(Annotated reads)
Cutoff . 3.56
Cutoff . 3.66
C
-Dox
+Dox
Top 25% expressed genes
Bottom 25% expressed genes
(DCM Reads / Sites)/ DCM depth
(CpG Reads / Sites)/ CpG depth
TSS
TES
D
Log10(CpG Unique Reads)
Log10(Mapped reads)
R² = 0.71
Log10(DCM Unique Reads)
E
DCM/CpG
F
Genes
DCM counts
G
-Dox DCM
+Dox DCM
H3K27ac
H3K4me3
H3K36me3
H3K27me3
H3K9me3
-4 Kb
4 Kb
H
-Dox CpG
+Dox CpG
I
RNA
Nanog
Dcx
J
Other
Pluripotency TFs
-log10(P-value)
Oct4-Sox2-Tcf-Nanog
Klf4
Sox2

**Figure S1. Benchmarking single cell state tracing using sciMeD-Seq and combinatorial indexing in mESCs.** (A) Single cell combinatorial indexing MeDseq (sciMeD-Seq) outline. Nuclei are isolated followed by fixation and nucleosome depletion. The gDNA is digested with LpnPI within intact nuclei. Barcoded adapters are ligated to digested gDNA and each nucleus is assigned to a specific barcode combination through several rounds of split-pool-ligation steps followed by P5-P7 indexing through library prep. (B) Density plot visualizing the distribution of annotated reads across all detected barcodes (n=~2000 nuclei were used for library preparation per condition). Cut-offs are based on Valley point analysis (cut-off: 3.56 $\log_{10}$ (n=1,399 nuclei) and 3.66 $\log_{10}$ transformed (n=1,939 nuclei)), with barcodes above this threshold representing nuclei retained for downstream analysis. (C) Metagene plots of pseudo-bulk single cell combinatorial indexing MeD-Seq (sciMeD-Seq). DCM methylation density (left panel) on the 25% highest expressed genes (solid line) and 25% lowest expressed genes (dashed line). Dox induced (blue) vs non-induced (red) mESCs are shown. Right panel shows the same features for CpG methylation density. (D) Number of unique CpG (left) and DCM (right) reads (y-axis) vs aligned reads (x-axis) per cell across conditions (±dox). Samples display a linear increase in unique read recovery as a function of sequencing depth (+dox: $R^2 = 0.71$). (E) Box plots showing percentage of aligned reads in filtered cells (top panel; median ~98.8%) and DCM/CpG ratio (bottom panel) for both non-induced (red) and induced (blue) cells (0.07 vs 0.5, median DCM/CpG ratio). The boxplot displays outliers (black dots), the median (black line), the interquartile range (box limits), and the whisker represents 1.5x of the interquartile range. (F) Scatter plot showing a linear relationship between unique DCM counts on genes (x-axis) and the number of genes recovered (y-axis), for both uninduced (red dots) and induced (blue dots) conditions. The dashed line indicates the threshold used to filter out cells with fewer than 300 detected genes. (G) Heatmaps showing scaled DCM read density across different enriched regions for histone modifications in mESCs: H3K27ac (7,603 regions), H3k4me3 (13,535 regions), H3K36me3 (85,125 regions), H3K27me3 (8,506 regions), and H3K9me3 (8,187 regions). Visualized is an 8kb region centered on the peak of the enriched regions. The top heatmaps represent a 24h dox induced sample, while the bottom heatmaps represent a non-induced sample. Boxes above the heatmap show collapsed metagene plots of the data. (H) Heatmaps of scaled CpG read density as shown in G. CpG reads were read density normalized. (I) Genome tracks of DCM read density for non-induced and induced sciMeD-Seq data covering the Nanog and Dcx genes. Bottom lane shows complementary RNA density from RNA-Seq data in mESCs [67]. RNA density is count per million normalized while DCM counts were adjusted for total reads. (J) Barplot visualizing the motifs enriched in DCM regions with signal in putative enhancers (EnhancerAtlas2.0). *Sox2*, *Klf4*, and *Oct4-Sox2-Tcf-Nanog* combinations are enriched across the top 100 motifs (FDR ~ 0)

Fig. S2

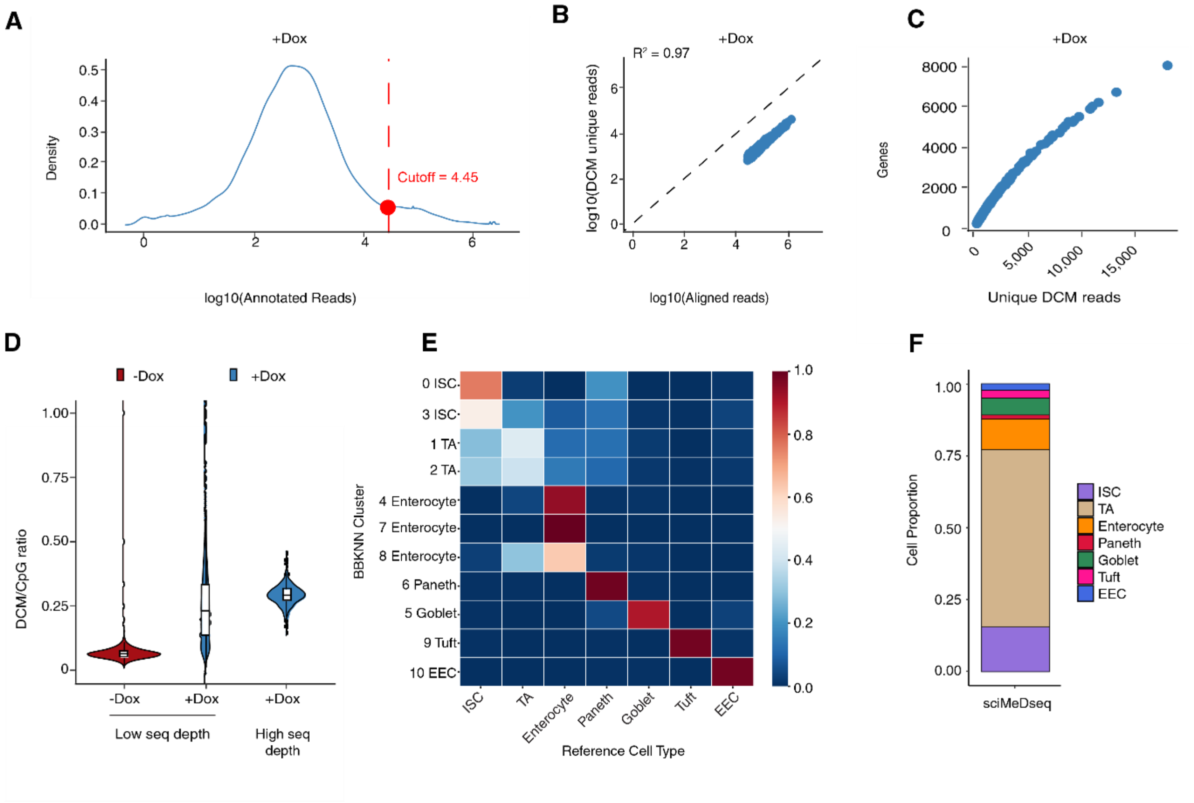
A
+Dox
Density
log10(Annotated Reads)
Cutoff = 4.45
B
+Dox
R² = 0.97
log10(DCM unique reads)
log10(Aligned reads)
C
+Dox
Genes
Unique DCM reads
D
-Dox
+Dox
DCM/CpG ratio
Low seq depth
High seq depth
E
BBKNN Cluster
0 ISC
3 ISC
1 TA
2 TA
4 Enterocyte
7 Enterocyte
8 Enterocyte
6 Paneth
5 Goblet
9 Tuft
10 EEC
ISC
TA
Enterocyte
Paneth
Goblet
Tuft
EEC
Reference Cell Type
F
Cell Proportion
sciMeDseq
ISC
TA
Enterocyte
Paneth
Goblet
Tuft
EEC

**Figure S2. Benchmarking single cell state tracing using combinatorial indexing in the mouse intestine.** (A) Density plot visualizing the number of annotated reads on a $\log_{10}$ scale (x-axis) across all detected barcodes (y-axis) in dox-induced enterocytes. The cut-off is visualized with the red dot and dashed line based on Valley point analysis (cut-off: 4.45 $\log_{10}$ transformed (n=516 nuclei vs expected barcodes ~500). (B) Graph showing the number of unique DCM reads recovered as a function of aligned reads per cell. A range of 623-41,734 DCM unique fragments were recovered across all cells. The relationship between sequencing depth and unique DCM recovery was highly consistent across filtered cells ($R^2$=0.97, log-log linear model). (C) Scatter plot showing the relationship between unique DCM counts (x-axis, mean=1,428 genes) and the number of genes per cell (y-axis; mean = 1,918 UMIs). nuclei with less than 200 detected genes were filtered out, resulting in a total of 516 nuclei for downstream analysis. (D) Violin plots showing the DCM/CpG ratio per cell across conditions. Samples were sequenced at low depth separately for both the non-induced (-Dox) and 48-hour induced (+Dox) conditions, with median ratios of 0.06 and 0.23, respectively to confirm induction prior to high depth sequencing. High depth sample (median DCM/CpG ratio = 0.29). The boxplot inside the violin plot displays, the median (black line), the interquartile range (box limits), and the whisker represents 1.5x of the interquartile range. (E) Heatmap showing the fraction of intersecting cells between Scanpy-derived clusters and RNA reference cell type labels. This comparison was used to assign cluster identities consistent with RNA reference annotations. (F) Cell proportion distribution for sciMeD-Seq dataset after BBKNN integration. Each color represents one cell type as shown.

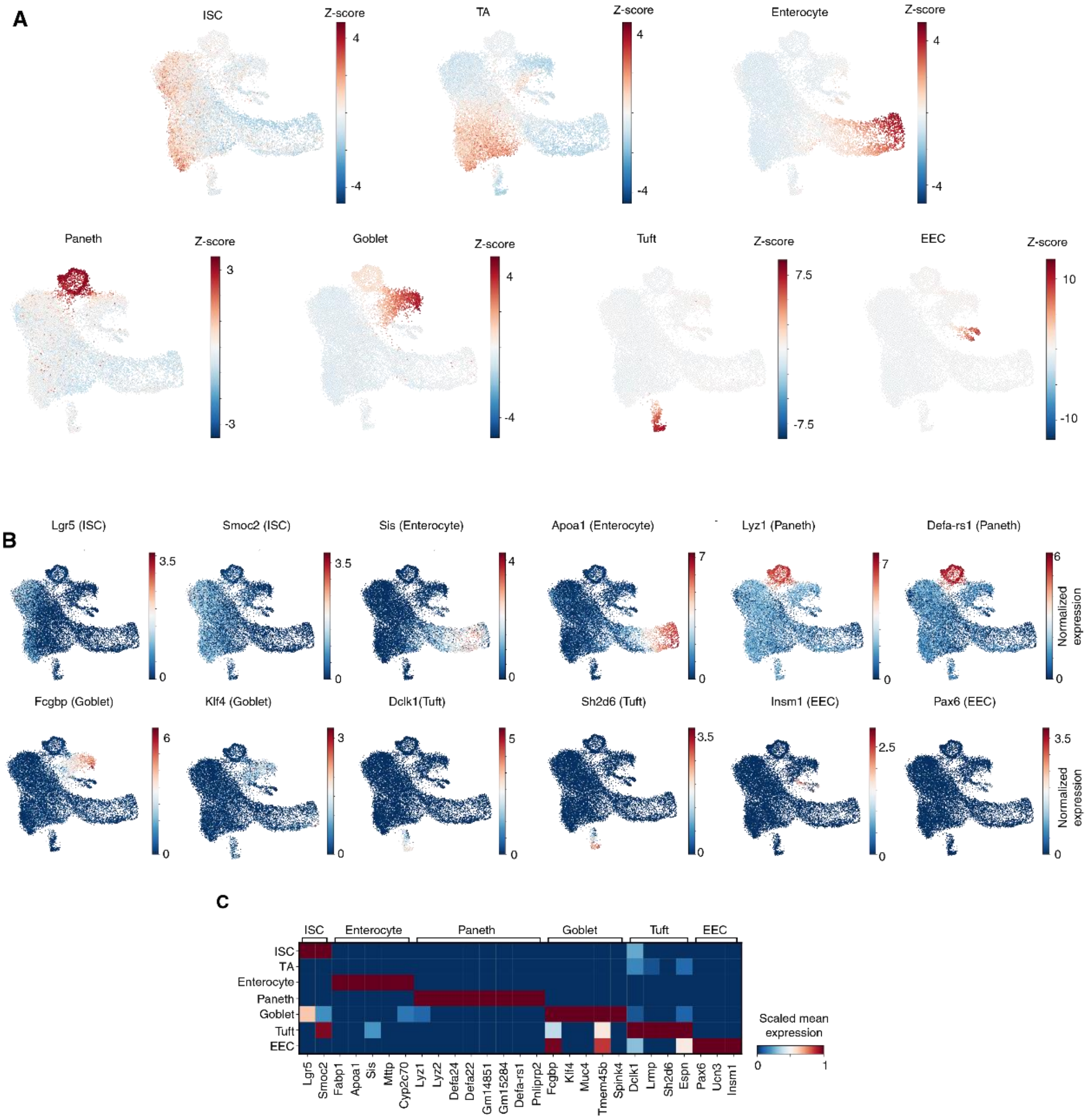
A
ISC
Z-score
TA
Enterocyte
Paneth
Goblet
Tuft
EEC
B
Lgr5 (ISC)
Smoc2 (ISC)
Sis (Enterocyte)
Apoa1 (Enterocyte)
Lyz1 (Paneth)
Defa-rs1 (Paneth)
Normalized expression
Fcgbp (Goblet)
Klf4 (Goblet)
Dclk1(Tuft)
Sh2d6 (Tuft)
Insm1 (EEC)
Pax6 (EEC)
C
ISC
Enterocyte
Paneth
Goblet
Tuft
EEC
Scaled mean expression
Lgr5
Smoc2
Fabp1
Apoa1
Sis
Mttp
Cyp2c70
Lyz1
Lyz2
Defa24
Defa22
Gm14851
Gm15284
Defa-rs1
Pnliprp2
Fcgbp
Klf4
Muc4
Tmem45b
Spink4
Dclk1
Lrmp
Sh2d6
Espn
Pax6
Ucn3
Insm1

**Figure S3. Z-scored modules for single cell states based on RNA and DCM data.** (A) UMAP visualizing the average gene set expression across cells for each cell type. The color indicates the scaled score levels for each module which is calculated by measuring the average normalized expression of each gene set subtracting by a background gene signal. Each panel indicates a separate cell type specific module score (shown for ISC (top left; n = 16 genes), TA (top middle; n = 14 genes), Enterocyte (top right; n = 17 genes), Paneth (bottom left; n= 27 genes), goblet (bottom 1-middle; n = 15 genes), tuft (bottom 2-middle; n=20 genes) and EEC (bottom right; n=18 genes) gene modules). (B) Log-transformed gene expression of cell-type-enriched markers visualized in UMAP space. Gene names appeared on top of UMAPs while in brackets the corresponding cell type marker is shown. (C) Matrix plot showing scaled normalized gene expression of cell type specific markers on DCM cells matching BBKNN integration labels and GMM assignments.

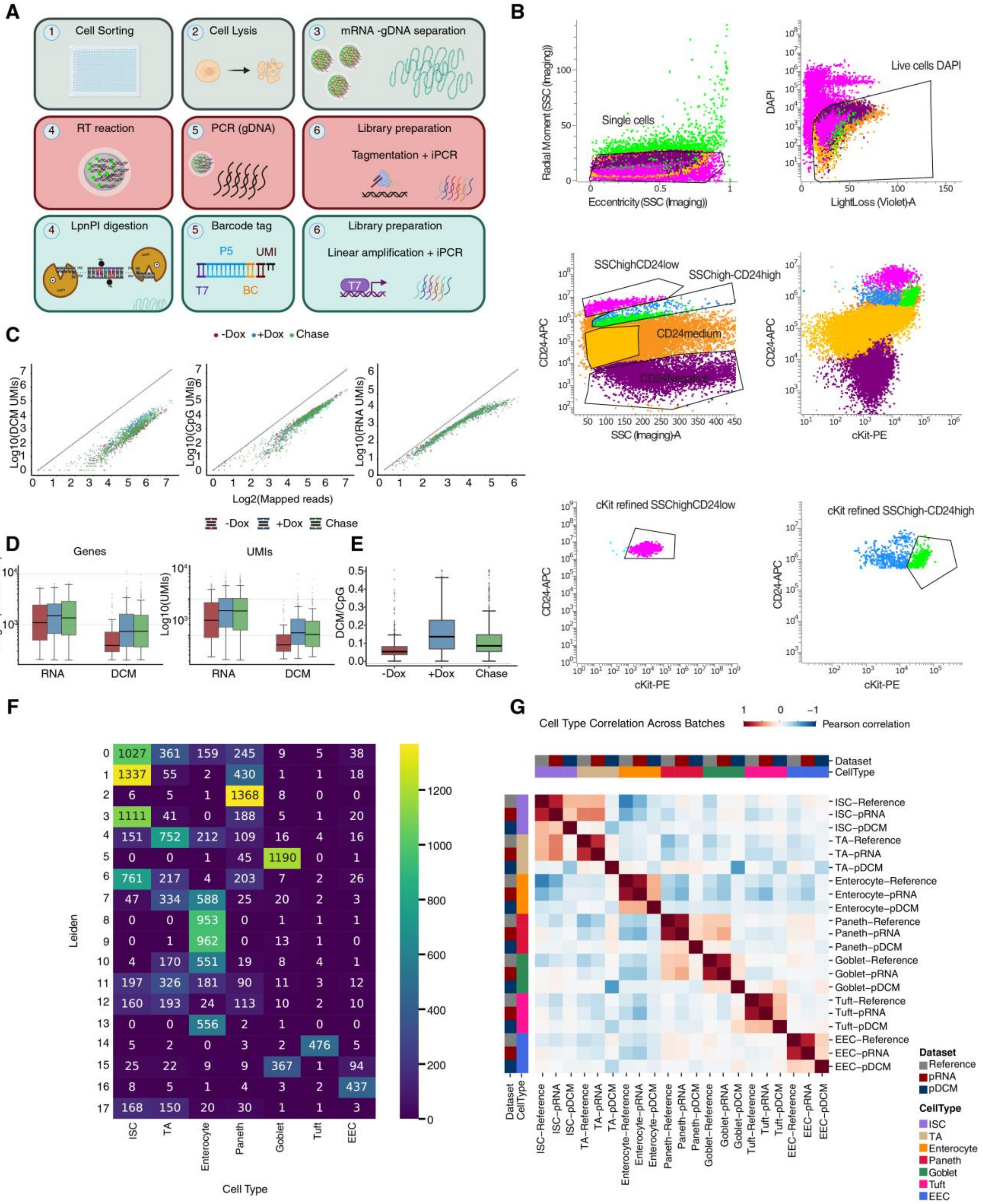
A
1 Cell Sorting
2 Cell Lysis
3 mRNA -gDNA separation
4 RT reaction
5 PCR (gDNA)
6 Library preparation
Tagmentation + iPCR
4 LpnPI digestion
5 Barcode tag
P5
UMI
T7
BC
6 Library preparation
Linear amplification + iPCR
T7
B
Radial Moment (SSC (Imaging))
Single cells
Eccentricity(SSC (Imaging))
DAPI
Live cells DAPI
LightLoss (Violet)-A
CD24-APC
SSChighCD24low
SSChigh-CD24high
CD24medium
CD24Negative
SSC (Imaging)-A
CD24-APC
cKit-PE
cKit refined SSChighCD24low
CD24-APC
cKit-PE
cKit refined SSChigh-CD24high
CD24-APC
cKit-PE
C
-Dox +Dox Chase
Log10(DCM UMIs)
Log10(CpG UMIs)
Log10(RNA UMIs)
Log2(Mapped reads)
-Dox +Dox Chase
D
Genes
Log10(# of Genes)
RNA DCM
UMIs
Log10(UMIs)
RNA DCM
E
DCM/CpG
-Dox +Dox Chase
F
Leiden
Cell Type
ISC TA Enterocyte Paneth Goblet Tuft EEC
0: 1027 361 159 245 9 5 38
1: 1337 55 2 430 1 1 18
2: 6 5 1 1368 8 0 0
3: 1111 41 0 188 5 1 20
4: 151 752 212 109 16 4 16
5: 0 0 1 45 1190 0 1
6: 761 217 4 203 7 2 26
7: 47 334 588 25 20 2 3
8: 0 0 953 0 1 1 1
9: 0 1 962 0 13 1 0
10: 4 170 551 19 8 4 1
11: 197 326 181 90 11 3 12
12: 160 193 24 113 10 2 10
13: 0 0 556 2 1 0 0
14: 5 2 0 3 2 476 5
15: 25 22 9 9 367 1 94
16: 8 5 1 4 3 2 437
17: 168 150 20 30 1 1 3
0 200 400 600 800 1000 1200
G
Cell Type Correlation Across Batches
1 0 −1 Pearson correlation
Dataset
CellType
ISC−Reference
ISC−pRNA
ISC−pDCM
TA−Reference
TA−pRNA
TA−pDCM
Enterocyte−Reference
Enterocyte−pRNA
Enterocyte−pDCM
Paneth−Reference
Paneth−pRNA
Paneth−pDCM
Goblet−Reference
Goblet−pRNA
Goblet−pDCM
Tuft−Reference
Tuft−pRNA
Tuft−pDCM
EEC−Reference
EEC−pRNA
EEC−pDCM
Dataset
Reference
pRNA
pDCM
CellType
ISC
TA
Enterocyte
Paneth
Goblet
Tuft
EEC

**Figure S4. QC metrics and benchmarking of scMeD&RNA-Seq.** (A) Schematic representation of the miniaturized scRNA&MeDseq protocol. (B) Gating strategy used to sort single live cells (top left and right panels) from various small intestine populations, originating from both crypt and villus compartments. $CD24_{Negative}$ (purple events, middle left and right panels), $CD24_{medium}$ (orange events, middle left and right panels), $CD24_{high}/SSC_{low}$ and c-$Kit_{high}$, (pink events, bottom left) and $CD24_{high}$ and c-$Kit_{high}$, green events, bottom right) were sorted. (C) Unique DCM (left panel), CpG (middle panel) and RNA (right panel) reads detected as a function of mapped read across cells for each condition (non-induced (-Dox), 48h dox (+Dox) and 48h dox followed by 72h chase (Chase)) as indicated. A range of 203- 690,631 CpG sites per barcode was captured for all conditions after filtering low quality cells, while a range of 200-26,828 DCM sites were captured for non-induced cells and 202- 110,138 DCM sites were detected for induced cells. Cells still show a linear increase in unique reads as a function of sequencing depth. (D) Box plots showing unique DCM and RNA counts and genes for each condition as indicated. Each box plot displays, the median (black line), the interquartile range (box limits), and the whisker represents 1.5x of the interquartile range. (E) Box plots showing DCM-to-CpG ratio indicating the induction across cells per condition (-Dox, +Dox (48h dox) and Chase (48h dox followed by 72h chase)). The boxplot displays outliers (black dots), the median (black line), the interquartile range (box limits), and the whisker represents 1.5x of the interquartile range. (F) Heatmap showing the number of intersecting cells between leiden-derived clusters and RNA reference cell type labels after integration of single cell datasets. This comparison was used to assign clusters identities consistent with RNA reference annotations. (G) Heatmap showing the Pearson correlation between depth-normalized expression profiles, followed by z-score scaling across datasets for each cell type: (Atlas and Large scRNA reference datasets (gray)[26], and scMeD&RNA-Seq (pRNA (red) and pDCM (blue)). Pearson correlation values are represented as a color gradient, with red indicating positive correlation and blue indicating negative correlation.

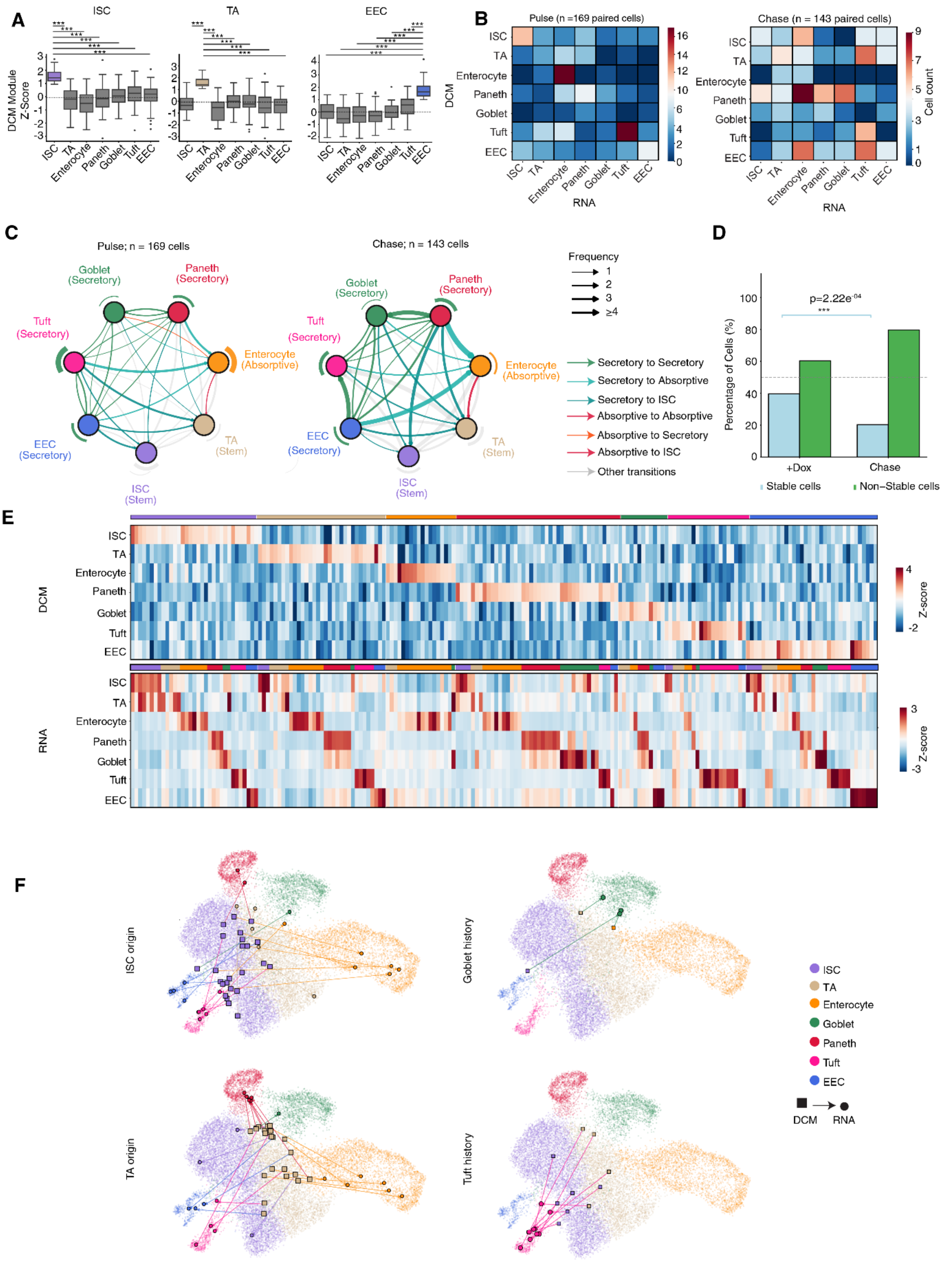

A
ISC
TA
EEC
DCM Module Z-Score
ISC TA Enterocyte Paneth Goblet Tuft EEC
B
Pulse (n =169 paired cells)
Chase (n = 143 paired cells)
DCM
RNA
Cell count
C
Pulse; n = 169 cells
Chase; n = 143 cells
Goblet (Secretory)
Paneth (Secretory)
Tuft (Secretory)
Enterocyte (Absorptive)
EEC (Secretory)
TA (Stem)
ISC (Stem)
Frequency
1
2
3
≥4
Secretory to Secretory
Secretory to Absorptive
Secretory to ISC
Absorptive to Absorptive
Absorptive to Secretory
Absorptive to ISC
Other transitions
D
p=2.22e-04
***
Percentage of Cells (%)
+Dox
Chase
Stable cells
Non–Stable cells
E
DCM
RNA
Z-score
F
ISC origin
Goblet history
TA origin
Tuft history
ISC
TA
Enterocyte
Goblet
Paneth
Tuft
EEC
DCM
RNA

**Figure S5. Cell transitions across secretory and absorptive lineages.** (A) DCM module Z-scores for marker gene sets specific to each cluster (cell type) calculated for indicated cell types following GMM-based cell assignments. Each panel indicates a separate cell type specific gene module tested (shown from left to right ISC, TA, and EEC gene modules). Colored boxplots highlight the cell type upon which module genes were selected and match the colors in F. Reference values are compared to other cell types using a pairwise t-test adjusted for multiple testing (FDR). Significance: *** ($p < 0.001$). (B) Heatmaps showing the number of cell state transitions between cell types under pulse (left) and chase (right) conditions based on GMM labelling for DCM cells and BBKNN integration labelling for RNA cells. Each heatmap depicts number of transitions between, cell types with DCM-based labelling representing past gene activity (x-axis) and RNA-based labelling representing current gene activity (y-axis). (C) Network plot depicting the transitions classified based on GMM for DCM and BBKNN integration for RNA. The left plot shows transitions within the pulse condition, while the right plot shows transitions within the chase condition. The nodes represent annotated cell types while edges transitions between them. Transitions are colored as indicted. Edge thickness indicates transition frequencies. (D) Percentage of confident cells (after GMM filtering) retaining or changing their state to another state per dox condition. Blue bars show cells that are stable (no transition) and green bars the cells switching to another state (One-sided binomial test between pulse (48h dox) and chase (48h dox / 72h chase) within stable cells, p=2.22e-04, respectively). (E) pDCM and pRNA data scaled across gene modules (gene sets representing a cluster or cell type) for each cell to highlight the relative contribution of each gene module. Z-score DCM and RNA module scores are calculated as the average depth-normalized read density of cluster specific gene markers (ISC, n=603; TA, n=295; Enterocyte, n=1,275; Paneth, n=123; Goblet, n=465; Tuft, n=696; EEC, n=475) for both pRNA and pDCM data, following by z-score scaling. The heatmaps focuses on filtered unilabelled cells transitioning between lineages as indicated in the annotated label colors above the heatmaps. (F) UMAP visualization of DCM cells assigned to multiple cell states (probability >0.85 for more than one label) and their corresponding transitions based on BBKNN integration. DCM cells are represented as rectangles colored according to their assigned labels, while RNA cells appear as circles.

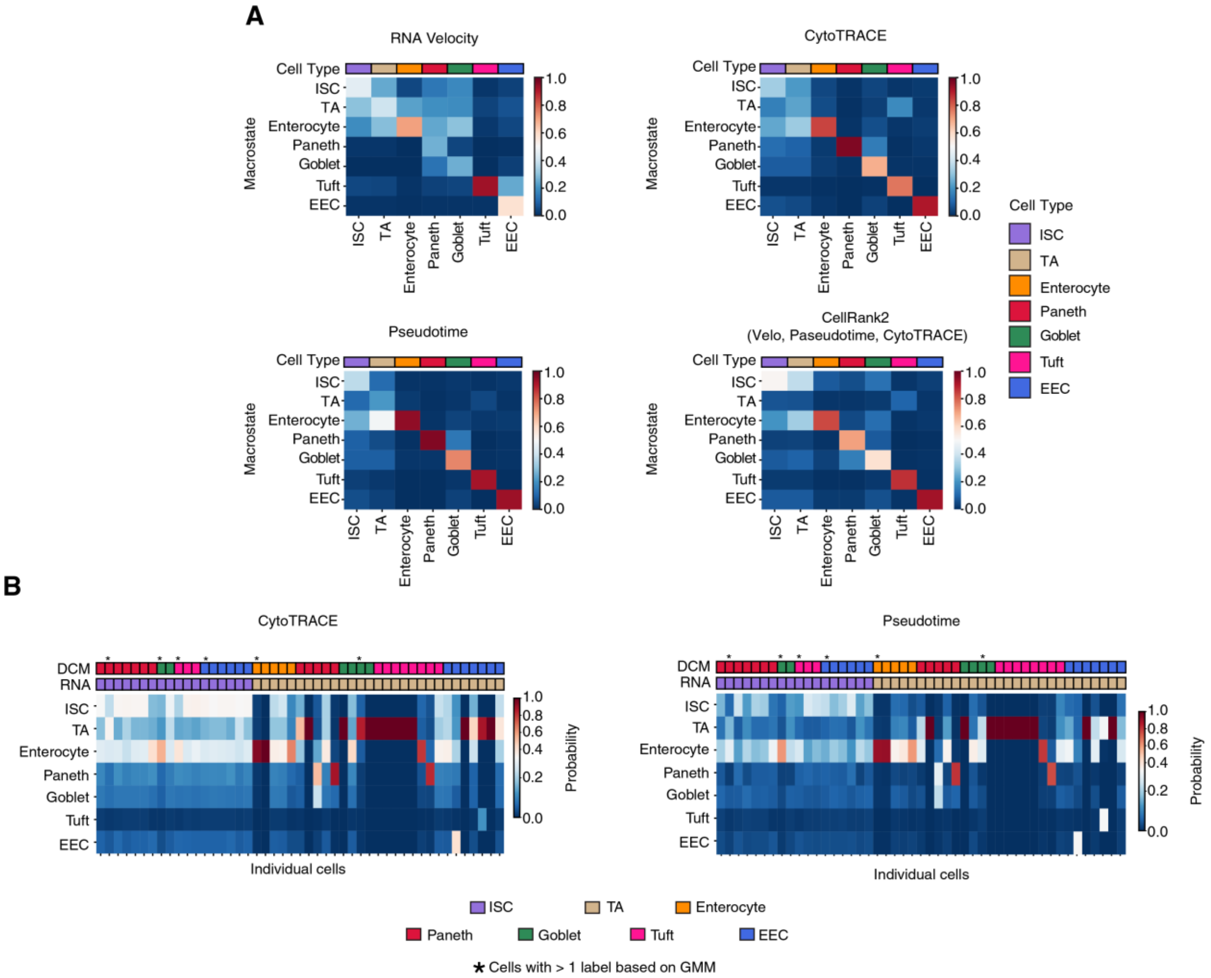
A
RNA Velocity
CytoTRACE
Pseudotime
CellRank2
(Velo, Paseudotime, CytoTRACE)
Cell Type
Macrostate
ISC
TA
Enterocyte
Paneth
Goblet
Tuft
EEC
1.0
0.8
0.6
0.4
0.2
0.0
B
CytoTRACE
Pseudotime
DCM
RNA
Individual cells
Probability
* Cells with > 1 label based on GMM

**Figure S6. Cell fate probabilities of secretory and absorptive cell types predicted by CellRank2.** (A) Heatmap displaying the probability of clusters to each macrostate, grouped by cluster identity. The probability is measured using RNA velocity (top left), CytoTRACE (top right), pseudotime (bottom left) or a combination of all three of them (bottom right). The corresponding cell type is labelled with their respective color above the heatmap. ISC and TA were set as the initial cluster while all cell types were considered terminal clusters, including ISCs, due to their self-renewal capacity. (B) Same analysis as in (A) but for individual cells that transition back to ISCs or Tas. Heatmaps include RNA velocity, CytoTRACE and pseudotime analysis as indicated. For each cell the GMM based DCM label (DCM) and RNA label (RNA) are indicated by colors. Hybrid states are assigned ('*'). Heatmaps display the probability of individual cells to transition towards each macrostate, grouped by cluster identity. The probability combines RNA velocity (30%), CytoTRACE (30%) and pseudotime (30%) modalities and cell connectivity (10%). Statistical significance is indicated as follows: *** ($p < 0.001$); ** ($p < 0.01$); * ($p < 0.05$); ns ($p > 0.05$).

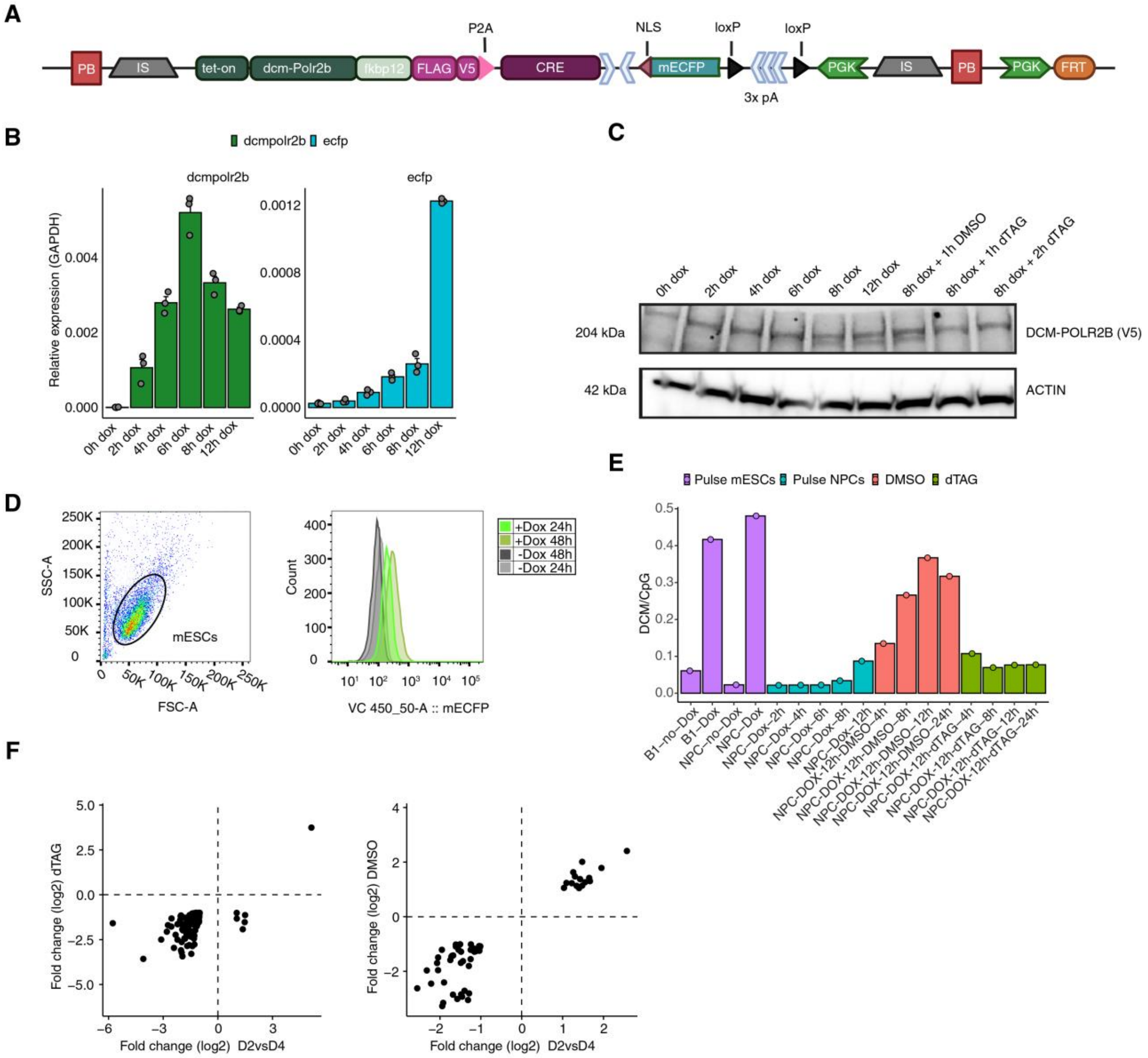

A
PB
IS
tet-on
dcm-Polr2b
fkbp12
FLAG
V5
P2A
CRE
NLS
mECFP
loxP
loxP
3x pA
PGK
IS
PB
PGK
FRT
B
dcmpolr2b
ecfp
Relative expression (GAPDH)
0h dox
2h dox
4h dox
6h dox
8h dox
12h dox
C
0h dox
2h dox
4h dox
6h dox
8h dox
12h dox
8h dox + 1h DMSO
8h dox + 1h dTAG
8h dox + 2h dTAG
204 kDa
DCM-POLR2B (V5)
42 kDa
ACTIN
D
SSC-A
FSC-A
mESCs
Count
+Dox 24h
+Dox 48h
-Dox 48h
-Dox 24h
VC 450_50-A :: mECFP
E
Pulse mESCs
Pulse NPCs
DMSO
dTAG
DCM/CpG
B1–no–Dox
B1–Dox
NPC–no–Dox
NPC–Dox
NPC–Dox–2h
NPC–Dox–4h
NPC–Dox–6h
NPC–Dox–8h
NPC–Dox–12h
NPC-DOX-12h-DMSO-4h
NPC-DOX-12h-DMSO-8h
NPC-DOX-12h-DMSO-12h
NPC-DOX-12h-DMSO-24h
NPC-DOX-12h-dTAG-4h
NPC-DOX-12h-dTAG-8h
NPC-DOX-12h-dTAG-12h
NPC-DOX-12h-dTAG-24h
F
Fold change (log2) dTAG
Fold change (log2) D2vsD4
Fold change (log2) DMSO
Fold change (log2) D2vsD4

**Figure S7. Benchmarking the dcm-polr2b-fkbp12 system in mESCs and NPCs.** (A) Schematic representation of the construct used to target mESCs for DCM cell tracing. Key components include a doxycycline-inducible promoter (Tet-On) driving expression of *dcm-polr2b*, fused to an FKBP12 degron for dTAG-mediated degradation. A FLAG-V5 epitope tag enables protein-level visualization. The construct also includes a Cre recombinase that excises a loxP-flanked stop cassette, enabling *mecfp* expression under a constitutive promoter to track induced cells. Insulator elements flank the construct to prevent integration based transcriptional effects, while PiggyBac and FRT sequences support both random and targeted (single-site) genomic integration. (B) *dcm-Polr2b* (left) and *mecfp* (right) relative RNA expression compared to *gapdh*. Dox induction was tested every 2h from 0 to 8h and 12h in mESCs$^{dcm\text{-}pol\text{-}fkbp12\text{-}cre}$. (C) Western blot showing DCM-POLR2B protein levels following induction. Protein lysates were extracted from dox induced mESCs every 2h from 0 to 8h and 12h followed by 500nM dTAG-13 or DMSO treatment as indicated. (D) Flow gates for mESCs after 24h and 48h dox induction. The first gate detects the cell population of interest while the second shows the eCFP expression across different conditions (dark and light gray: -Dox; light green: 24h +Dox; dark green 48h +Dox). (E) Bar graph showing DCM/CpG ratio based on Med-Seq across different induction times for samples in mESCs$^{dcm\text{-}pol\text{-}fkbp12\text{-}cre}$ (B1) and NPCs. The Positive controls (B1-Dox and NPC-Dox induced for 24h Dox) are labelled in purple together with their negative controls (B1-no-Dox and NPC-no-Dox). The pulse cells are labelled in blue and showing induction times as indicated DMSO or dTAG-13 treatment (red and green bars, respectively: shown for treatment time in the presence of Dox). (F) $Log_2$ fold-change of differentially DCM methylated genes. The comparisons were performed between early neural differentiation of D2 vs D4 following a dox pulse (x-axis) and D2 dox pulse followed by 24h dTAG (left panel) or DMSO (right panel) treatment vs D4 dox induction (y-axis). Genes with FDR $< 0.05$ were selected between D2 vs D4 pulse comparison.

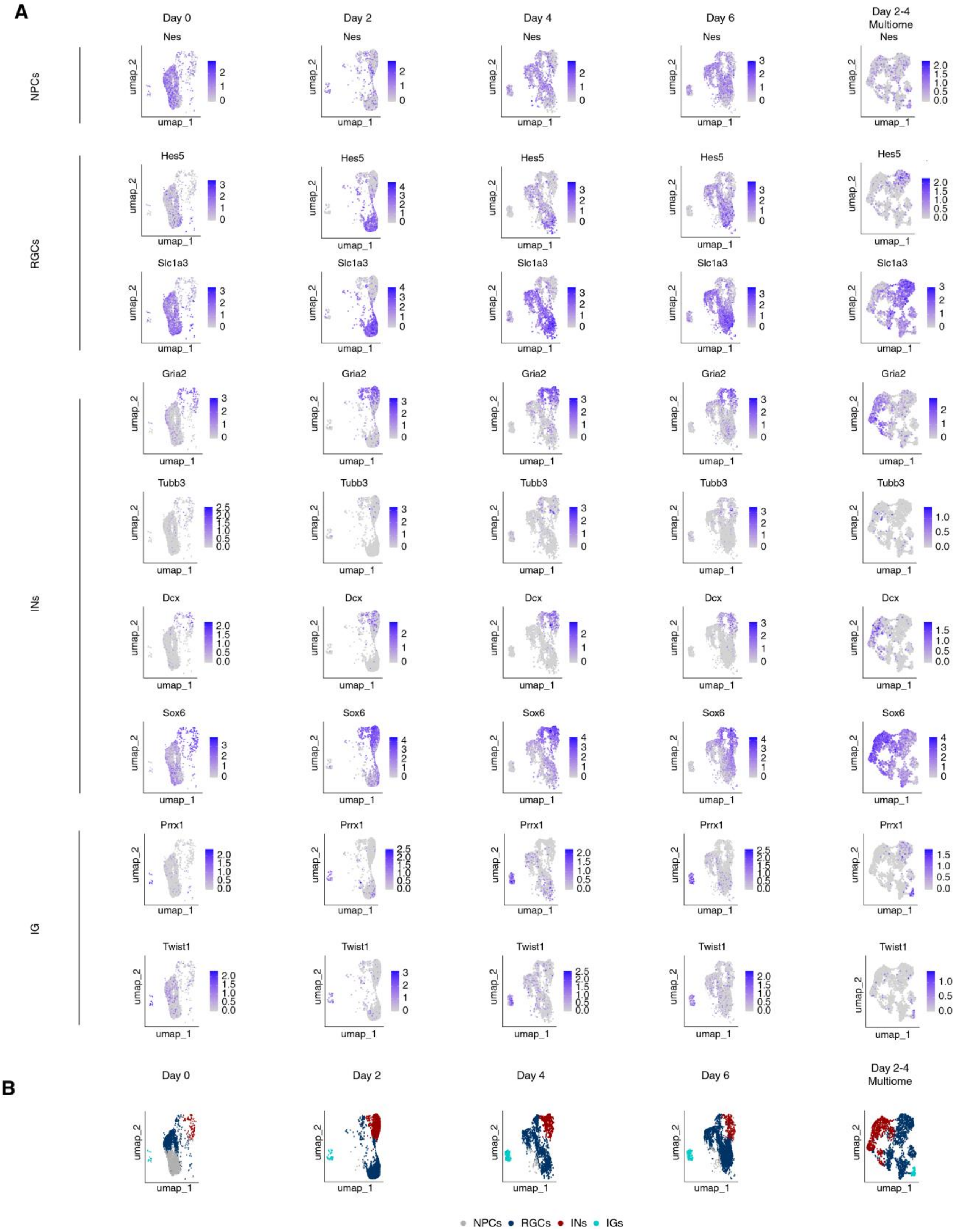

A
Day 0
Day 2
Day 4
Day 6
Day 2-4 Multiome
NPCs
Nes
RGCs
Hes5
Slc1a3
INs
Gria2
Tubb3
Dcx
Sox6
IG
Prrx1
Twist1
umap_1
umap_2
B
Day 0
Day 2
Day 4
Day 6
Day 2-4 Multiome
NPCs
RGCs
INs
IGs

**Figure S8. Cluster gene markers for NPCs, RGCs, INs and IGs.** (A) UMAP visualizations showing gene expression of key marker genes across neural differentiation for scRNA datasets at days 0, 2, 4, 6, as well as multiome data (D2-D4 multiome). Gene expression is color-mapped across cells, with intensity indicating expression level. Marker genes include mESCs$^{dcm\text{-}pol\text{-}fkbp12\text{-}cre}$ derived neural progenitor cells (NPCs$^{dcm\text{-}pol\text{-}fkbp12\text{-}cre}$, *Nes*), radial glial cells (RGCs; *Hes5*, *Slc1a3*), immature neurons (INs; *Gria2*, *Tubb3*, *Dcx*, *Sox6*), and immature glial differentiation (IG; *Prrx1*, *Twist1*). (B) UMAP visualizations of cell clusters per timepoint and dataset.

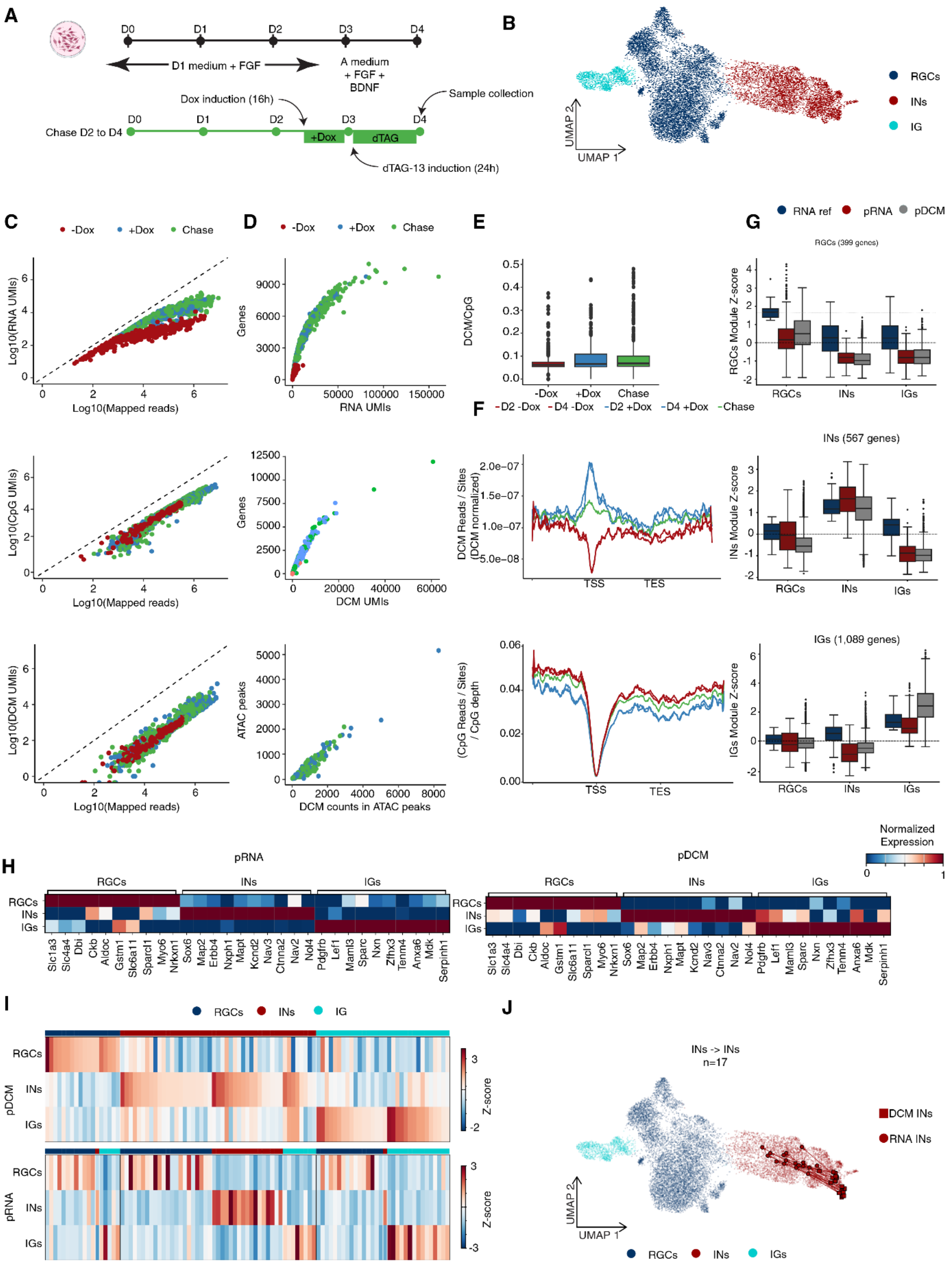

A
D0
D1
D2
D3
D4
D1 medium + FGF
A medium + FGF + BDNF
Dox induction (16h)
Sample collection
Chase D2 to D4
+Dox
dTAG
dTAG-13 induction (24h)
B
UMAP 2
UMAP 1
RGCs
INs
IG
C
-Dox
+Dox
Chase
Log10(RNA UMIs)
Log10(Mapped reads)
Log10(CpG UMIs)
Log10(DCM UMIs)
D
Genes
RNA UMIs
DCM UMIs
ATAC peaks
DCM counts in ATAC peaks
E
DCM/CpG
F
D2 -Dox
D4 -Dox
D2 +Dox
D4 +Dox
DCM Reads / Sites (DCM normalized)
(CpG Reads / Sites) / CpG depth
TSS
TES
G
RNA ref
pRNA
pDCM
RGCs (399 genes)
RGCs Module Z-score
INs (567 genes)
INs Module Z-score
IGs (1,089 genes)
IGs Module Z-score
H
pRNA
pDCM
Normalized Expression
Slc1a3
Slc4a4
Dbi
Ckb
Aldoc
Gstm1
Slc6a11
Sparcl1
Myo6
Nrkxn1
Sox6
Map2
Erbb4
Nxph1
Mapt
Kcnd2
Nav3
Ctnna2
Nav2
Nol4
Pdgfrb
Lef1
Maml3
Sparc
Nxn
Zfhx3
Tenm4
Anxa6
Mdk
Serpinh1
I
Z-score
J
INs -> INs
n=17
DCM INs
RNA INs
IGs

**Figure S9. Benchmarking and QC metrics, for neural differentiation.** (A) Schematic representation of neural differentiation. The black timeline shows the differentiation protocol from day 0 to day 4 (D0-D4), while the green timeline represents the chase. Black arrows indicate dox induction, dTAG destruction and sample collection time points. (B) UMAP visualizations of BBKNN integrated data from scRNA-Seq day 2 (D2) (n=2,535), scRNA-Seq day 4 (D4) (n= 2,315), and multiome (D2–D4 mix; n = 9,433) datasets, combined with scMeD&RNA-Seq data (scDCM (n = 367) and scRNA (n = 727) post filtering). Clusters are annotated representing specific cell states and visualized in UMAP space. (C) Graphs showing quality control metrics for scMeD&RNA-Seq. Shown are UMI counts as a function of mapped reads per cell (x-axis) across induction conditions as indicated. (D) Graphs showing number of genes identified as a function of RNA UMIs (top graph) or DCM UMIs (middle graph) and number of ATAC peaks (bottom graph). (E) Boxplots showing DCM/CpG ratio for samples as indicated. The boxplot displays outliers (black dots), the median (black line), the interquartile range (box limits), and the whisker represents 1.5x of the interquartile range. (F) Metagene plots of the top 25% most highly expressed genes, based on reference 10x datasets (Days 0-6). DCM signal is shown for induced and non-induced day 2 and day 4 samples (blue / red), as well as chase samples (green), normalized by DCM site coverage and DCM depth. The bottom panel shows CpG signal across the same gene set. (G) Z-module scores for cells based on the marker gene sets corresponding to each cluster (cell type) as indicated. Shown is RNA reference data (blue boxplot), pRNA data (red boxplot) and pDCM data (grey boxplot). Z-scores for each cluster specific marker gene set (RGCs, n=399; IN, n=567; and IG, n=1,089), indicate how strongly the cell's gene expression pattern deviates from the average expression pattern across all cells in the dataset. (H) Top gene markers per cluster for pRNA (left panel) and pDCM (right panel) data. Color indicates scaled, depth-normalized gene expression levels. (I) Heatmaps showing Z-score DCM and RNA module scores calculated as the average depth-normalized read density of cluster specific gene markers (RGCs, n=399; IN, 567; and IG, n=1,089) for both pRNA and pDCM data, following by z-score scaling. Top and bottom panel shows scores for DCM and RNA data as shown. The heatmap focuses on filtered cells transitioning between RGCs, INs and IGs lineages based on Harmony data integration. (J) UMAPs showing the transitions from filtered INs to Ins in UMAP space. Arrows indicate the direction of transition between paired DCM (square) and RNA (circle) profiles of individual cells. Cells are colored by their respective cell types.

**Table S1.** Selected literature gene markers and top sensitive genes using multinomial logistic regression (Coefficient >0.5) and identified unique genes per cell type based on differential expression analysis, using a t-test (log2FC > 1, FDR < 0.0001).

**Table S2.** Oligo sequences needed for sciMeDseq protocol and library preparation.

**Table S3.** Oligo sequences needed for the scMeD&RNAseq protocol and library preparation.

**Table S4.** Oligo design for PCR, qPCR and CRISPR/Cas.
